# Secondary metabolites and the antimicrobial potential of five different *Coleus* species in response to salinity stress

**DOI:** 10.1101/220368

**Authors:** Divya Kotagiri, Shaik Khasim Beebi, Kolluru Viswanatha Chaitanya

## Abstract

Salinity is one of the major abiotic stresses that affects the growth and productivity of plants. The presence of soluble salts at high concentration near the root system restricts the uptake of water by plants. Plants grown under saline conditions possess higher amounts of secondary metabolites compared with those grown under normal conditions. The use of traditional medicine to treat infectious diseases is increasing day by day throughout the world. Developing novel drugs with antimicrobial potential from the source of medicinal plants is receiving attention to replace the use of synthetic drugs and to combat the growth of multi-drug resistant strains. Thus screening of medicinal plant extracts is carried out to evaluate their antimicrobial potency. The present study aimed at determining the secondary metabolites and antimicrobial potential of leaf, stem and root ethanol and chloroform extracts of five different *Coleus* species; *C.aromaticus, C.amboinicus, Cbarbatus, C.forskohlii* and *C.zeylanicus* subjected to salinity stress. The up regulation in the content of plant bioactive compounds along with the antimicrobial activities of ethanol and chloroform extracts under the influence of salinity stress have been observed during the study in *Coleus.* The leaf, stem and root extracts of all the five *Coleus* species showed good antimicrobial activity against the tested pathogenic strains. The leaf extracts of *Coleus* showed higher inhibitory activity compared to the stem and root extracts. Ethanol extracts showed higher anti-microbial activity ranging from 1.5-100 mg/ml compared with the chloroform extracts ranging from 0.97-250 mg/ml respectively. The study revealed that the increased antimicrobial activity with increasing salinity might be due to the up regulation of secondary metabolites. The leaf, stem and root extracts of *Coleus* showed effective antimicrobial activity against the pathogenic strains even under saline conditions is due to the up regulation of secondary metabolites which provides a scope of developing novel drugs to treat infectious diseases.

## Introduction

Infections caused by various bacterial and fungal pathogens are becoming a major threat to public health of the growing population in the developing countries. Usage of improper and synthetic medicines, mismanagement and maladministration of antibiotics along with the microbial mutations is leading to the development of multi-drug resistant pathogenic strains along with the side effects, enabling to search for the novel compounds with resistance to the emerging new strains (Thuy et al. 2016). Apart from the misuse of antibiotics, multi-drug resistant strains acquire resistance by several mechanisms like target site modification, metabolic inactivation and the efflux pumps expression leading to the antibiotic efflux (Yala et al. 2001; Hooper 2001). The emergence of new pathogens accounting for many infectious diseases along with the antibiotic resistance and increasing failure of chemotherapy is the largest causes of death in tropical countries. Unavailability of vaccines for most of these diseases enables the discovery of novel natural antibacterial agents for efficient treatment against infectious diseases. Plants synthesize a variety of secondary metabolites with potential anti-inflammatory, antimicrobial and antioxidant properties. The different parts of the plant like leaf, root, stem, flower, fruit, twigs etc. can be used as antimicrobial agents due to the presence of secondary metabolites (Seyyednejad et al. 2010). Secondary metabolites such as flavonoids, alkaloids, tannins and phenolic compounds provide protection against bacteria, fungi, viruses and insects used for the discovery and the development of novel drugs (Ghazghazi et al. 2015). The search of the plants with the efficient antioxidative defense system as well as capable of producing secondary metabolites with strong antimicrobial properties is being received much attention as a replacement for synthetic drugs. Since ancient times, plants with effective medicinal values have been used as the promising sources for the treatment of various ailments due to the presence of phytochemicals with therapeutic properties, which are the cheapest and safe alternative sources (Odeja et al. 2015).

Plants are frequently subjected to a variety of harsh environmental stresses such as scarcity of water, extreme temperatures, high soil salinity, herbivore attack, and pathogen infection diminishing their productivity (Sewelam et al. 2016). Salinity refers to the presence of different salts like sodium chloride, calcium sulphates, magnesium and bicarbonates in water and soil (Ouda 2008). Due to excessive use of fertilizers, irrigation with low quality water and desertification, cultivated soils are getting more saline worldwide (Ramadoss et al. 2013). Soil lands with high level of salt concentrations induces physiological and metabolic changes in plants affecting their seed germination, growth, development, yield and also decreases the rate of respiration and photosynthesis in plants. The uptake of water and absorption of essential nutrients by plants is restricted due to the presence of soluble salts exerting high osmotic pressure which ultimately affects the growth of plants (Tester and Devenport 2003). In addition to the growth and yield, the composition of bioactive compounds present in the aromatic and medicinal plants is affected by salinity (Gil et al. 2002). The increased levels of plant secondary metabolites such as phenols, flavonoids, tannins, alkaloids etc... under the influence of increased salt concentrations as a part of defence mechanism have been reported (Kate 2008). The preliminary screening of phytochemicals gives an idea about the type of compounds produced by plants and their quantification both under normal and saline conditions will be useful in extracting the compounds of interest in pure form followed by the identification of those metabolites in order to detect their significance in human health.

Medicinal plants are good sources of various secondary metabolites belong to the class of natural anti-oxidants useful in curing many diseases and as free radical scavengers (Wong et al. 2006, Adom et al. 2005). The presence of bioactive compounds is mainly responsible for anti-inflammatory and antioxidant properties of medicinal plants can be used as potential chemo preventives. Secondary metabolites or plant bioactive compounds are low molecular weight compounds distributed largely in plants play a major role in the adaptation of plants to different environmental changes and in overcoming stress constraints also used in neutralizing free radicals. The colour, smell, flavour and the defence mechanism against pathogens in plants is due to the presence of phytochemicals (Aziagba et al. 2017). The phenolic components such as flavonoids, phenolic acids and phenolic diterpenes are mainly responsible for antioxidative activity in medicinal plants due to redox properties involved in neutralizing free radicals, decomposing peroxides, quenching singlet and triplet oxygen (Lee et al. 2004; Ksouri et al. 2007). The concentration of bio-active compounds produced by plants depends mainly upon the growth conditions and especially under stress conditions influence the metabolic pathways leads to the accumulation of related natural compounds possess activity to scavenge reactive oxygen species (ROS). The common response observed in salt-stressed plants are the generation of ROS, highly reactive responsible in damaging cell structures, nucleic acids, lipids and proteins (Vaidyanathan et al. 2003). Plants possess medicinal value with anti-inflammatory and antimicrobial activities; acquire resistance to stress induced ROS is due to the presence of several bio-active compounds (Foyer et al. 1994).

The presence of phenolic compounds in medicinal plants is responsible for antimicrobial, anti-inflammatory, anti-thrombotic, vasodilatory, cardio protective and anti-allergic properties (Balasundram et al. 2006). The synthesis and accumulation of polyphenols are stimulated in response to salinity stress resulting in considerable variations in their quantity and quality. Flavonoids are one of the important classes of plant secondary metabolites protects plants from harmful UV rays and also from herbivores capable of transferring electrons to free radicals and to chelate and activate the enzymes with anti-oxidant properties inhibits free radical producing enzymes. The biological properties such as anti-viral, anti-malarial and the cholesterol synthesis inhibition are due to the presence of terpenoids (Indumathi et al. 2014). Thus the salt stressed medicinal plants can be used for economic purposes as they are a potential source of bioactive compounds (Valifard et al. 2014). The major phytoconstituents of *Coleus* reported so far are flavonoids, glycosides, phenolic and volatile compounds, but the quantitative analysis of secondary metabolites during salinity stress are less explored.

The presence of bio-active compounds in the leaf, stem and root extracts of *Coleus* possessing the property of antimicrobial activity have potential to damage ROS and the activity of free radicals, helps to maintain proper health by combating infectious diseases. The presence of ROS can react readily and oxidize various biomolecules like lipids, carbohydrates, DNA and proteins, mainly responsible for the human diseases such as ulcers, inflammation, autoimmune disorders and viral infections (Surh and Ferguson 2003). Medicinal plants are used in many countries to treat diseases as they are rich sources of compounds possessing antimicrobial property. More than 80% of world population depend on traditional medicine for their health care needs reported by WHO (World health organization) (Malleswari et al. 2017). Depending upon the type of solvent used, plant extracts can be administered to the patients either as raw or tisanes, nebulisate and as tinctures. Medicinal plants with secondary metabolites are capable of inducing specific physiological actions on the human body (Joshi and Parle 2006) and are a source of antioxidant (Nahak and Sahu 2010; Pandey and Madhuri 2010) and antimicrobial compounds (Maragathavalli et al. 2012; Sharma et al. 2012).

Genus *Coleus* is a perennial branched aromatic herb that belongs to the family of *“Lamiaceae”* can be grown indoor as well as outdoor possesses biological activity against various infectious diseases and a number of pharmacological effects. Five *Coleus* species considered for the study and cultivated are *C.aromaticus, C.amboinicus, C.forskohlii, C.barbatus* and *C.zeylanicus. Coleus aromaticus* possess antioxidant and anti-microbial properties and the leaves are used to treat cholera, diarrhoea, malarial fever, halitosis, convulsions, epilepsy, asthma, cough, flatulence, bronchitis, hepatopathy, anorexia, cephalagia, otalgia, dyspepsia, colic, hiccough, and strangury (Warrier et al. 1995). *Coleus forskohlii* is an aromatic herb grown under tropical to temperate conditions produces diterpenoid from its tuberous root called forskolin. It is used to treat painful urination, hypertension, insomnia, convulsions, eczema, respiratory disorders and congestive heart failure. It also possesses therapeutic features of curing asthma, psoriasis and cancer. Forskolin is used to prevent blood clotting helps in nerve regeneration, activates adenylate cyclase enzyme and to reduce the intraocular pressure in glaucoma. The root extracts of *Coleus forskohlii is* used to treat eczema and skin infections, also used to kill worms in the stomach. *C.forskohlii* used widely for curing several disorders like intestinal disorders, respiratory disorders, heart diseases, asthma, bronchitis, convulsions, insomnia, burning sensation, epilepsy and constipation (Ammon and Muller 1985). *C.forskohlii* is found to be effective in treating obesity, congestive heart failure, hypertension, psoriasis, glaucoma, asthma, depression and cancer metastasis. Apart from the medicinal value of this plant, *forskohlii* also contains essential oils used in the food industries as flavouring agents and as an anti-microbial compound (Chowdhary and Sharma 1998). *C.amboinicus* is considered as carminative, lactagogue, analgesic, anti-septic and anti-pyretic. The leaf extracts of *C.amboinicus* is used to treat headache, toothache, bites, burns and also effective against malaria parasite. *C.barbatus* is a perennial, succulent branched fleshy herb grows up to the height of 15-40 cm between 1000-2600 m altitudes above sea level used as a stimulant in the treatment of cough. The aerial parts of the plant have cytotoxic, anti-tumour and diuretic activities, also used in the treatment of gums and teeth disorders. The major active compounds present in this plant were diterpenes, triterpenes, tormentic acid, α‐ amyrin and the flavones 3,7 dimethyl quercetin, sitosterol and kumatakinin. *Coleus zeylanicus* has astringent and stomachic properties used in the treatment of fever, common cold, asthma, dysentery, diarrhoea, vomiting, burning sensation, small pox, eye diseases, worm diseases, chronic ulcers, dental diseases and thirst. The different parts of the plant like leaf, root and stem are rich in medicinal value. In the present study, efforts have been made to evaluate the antimicrobial potential of *Coleus* leaf, root and stem ethanol and chloroform extracts under normal and saline conditions.

## Materials and methods

### Coleus plants & salinity stress treatment

Five *Coleus* species, *aromaticus, amboinicus, zeylanicus, forskohlii* and *barbatus* were propagated in the GITAM University botanical garden in 12 inch pots under 720 minutes natural photoperiod [Irradiance (400-700 nm) of 1600-1800 µ mols m^-2^ s^-1^] with day/night temperatures of 30°C/23°C with an approximate air humidity of 60%. The pots were arranged in rows 1 m apart and the plants were irrigated daily. Three months old plants with uniform growth were selected for this study. *Coleus* plants of all varieties were then separated into four groups, namely control (0), mild (100 mM), moderate (200 mM) and severe (300 mM). Control plants were watered daily and salt-stressed plants were treated with 250 ml of 100, 200 and 300 mM Nacl solutions twice a day for a period of 1 week. Third or fourth leaf from the top of the plant was collected for all the experiments.

## Quantitative estimation of secondary metabolites

### Estimation of Phenols

Phenols estimated spectrophotometrically using Folin-Ciocalteau reagent which gives a blue colour complex measured at 650 nm. 0.5 g tissue was homogenized in 80% ethanol and centrifuged for 20 min at 10,000×g. The extracts were pooled together after repeated extraction with 80% ethanol and allowed to dry. The residue obtained was dissolved in 5 ml of distilled water. 2 ml of the aliquot was made up to 5 ml with distilled water and 0.5 ml of 1N Folin-Ciocalteau reagent and 2 ml of 20% Na_2_CO_3_ were added and incubated in a boiling water bath exactly for 1 minute. After cooling, absorbance of the samples was measured at 650 nm (Malick and Singh 1980).

### Estimation of Flavonoids

Flavonoids were estimated according to Chang et al. 2002. 0.5 grams of plant material were added to 5 ml of 8% methanol and extracted for 48 h by shaking at room temperature and centrifugation at 10,000×g for 20 min. To 0.5 ml of extract, 1.5 ml of methanol, 0.1 ml of 10% aluminium chloride, 0.1 ml of 1 M potassium acetate and 2.8 ml of distilled water were added and incubated at room temperature for 30 minutes. Absorbance of the samples measured at the wavelength of 415 nm.

### Estimation of Tannins

Tannins were estimated according to Polshettiwar et al. 2007 by the addition of 0.5 g *Coleus* tissue to 25 ml of distilled water and incubated at 100°C for 30 minutes, centrifuged at 10,000×g for 20 min. To 1 ml of extract, 1 ml of Folin-Denis reagent and 2 ml of sodium carbonate solutions were added and the volume was made up to 5 ml with distilled water, incubated at room temperature for 30 minutes and the absorbance was measured at 700 nm. Tannic acid was used as a standard for the preparation of calibration curve.

### Estimation of Anthraquinones

Anthraquinone content in the *Coleus* was determined by adding 0.05 g of dried tissue in 50 ml of distilled water extracted by shaking for 16 h. The contents were incubated at 70°C and 50 ml of 50% methanol was added and filtered. Absorbance of the filtrate was measured at 450 nm. Calibration standards were prepared using alizarin and purpurin at a concentration of 0.01 mg per 1 ml (Soladoye and Chukwuma 2012).

### Estimation of Alkaloids

Alkaloid content was estimated by adding 100 mg of dried *Coleus* tissue to 40 ml of 95% ethanol refluxed for about half an hour and then filtered. The volume of the filtrate was adjusted to 50 ml with 95%) ethanol and subjected to evaporation. The residue obtained was treated with 3 ml of 1N Hcl and allowed to stand for 2 h hydrolysis. 3 ml of 1N NaOH was added, followed by the 2 ml concentrated acetic acid and the volume being adjusted to 10 ml with distilled water. 1 ml of this solution was made up to 5 ml with 20% acetic acid and added to 5 ml of acetate buffer, 1 ml of 0.05% methyl orange and 5 ml chloroform. After few minutes chloroform layers are withdrawn, added with a pinch of Na_2_SO_4_ and the absorbance was measured at 420 nm. Solasodine was used as standard for calibration (Muthumani et al. 2010).

### Estimation of Terpenes

Terpenes were estimated spectrophotometrically by adding 10 ml of petroleum ether to 1 gram of leaf, stem and root powder and extracted with shaking for 15 min. The extract was filtered and the absorbance was measured at 420 nm (Mboso et al. 2013).

### Estimation of Steroids

Estimation of steroids was done by adding 2 ml of 4N H_2_SO_4_, 2 ml of 0.5% FeCl_3_ and 0.5 ml of 0.5% potassium hexa cyanoferrate to 1 ml of methanolic extract. The contents were incubated at 70°C for 30 min, allowed to cool and made up to the volume of 10 ml with distilled water. Absorbance of the samples was measured at 780 nm (Narendra et al. 2013).

### Estimation of Saponins

Saponins were estimated according to Brunner 1984. 1 g of fine powdered sample was weighed accurately and added to 100 ml of isobutyl alcohol and extracted with shaking for 5 h and then filtered. To the filtrate, 20 ml of 40% saturated magnesium carbonate solution was added and again subjected to filtration through a filter paper; a clear colourless filtrate was obtained. To 1 ml of the filtrate, 2 ml of 5% FeCl_3_ solution was added and the volume made up to 50 ml with distilled water. The contents were allowed to stand for 30 minutes at room temperature to develop a deep red colour and the absorbance of the samples was measured at 380 nm. A calibration curve was prepared using dioxgenin concentrations ranging from 0-100 μg.

### Estimation of Cardiac glycosides

1 g of *Coleus* tissue powder was added to 10 ml of 70% alcohol and extracted for 2-3 h followed by the filtration. 4 ml of the filtrate was added to 5 ml of 12.5% lead acetate and the volume made up to 50 ml with distilled water. The solution was again filtered and 5 ml of 4.77% disodium hydrogen orthophosphate was added to 25 ml of filtrate resulting in the formation of precipitate removed by a third round of filtration. A 5 ml of freshly prepared Buljet’s reagent was added to 5 ml of clear solution obtained after filtration and incubated at room temperature for 1 h. The absorbance of the samples was measured at 595 nm and the calibration curve was prepared using 0.02% Digitoxin dissolved in chloroform-methanol at the ratio of 1:1 v/v (El-olemy et al. 1994).

### Estimation of Lignins

Estimation of lignins was done by weighing 100 mg of dry *Coleus* tissue and 1 ml of 72% sulphuric acid was added, incubated at 30°C for 1h with occasional stirring. 28 ml of distilled water was added and the beaker was incubated at 120°C for 1 h. The contents were filtered and the residue obtained after filtration was dried overnight at 105°C and determined the weight (AIR) whereas the filtrate obtained measured at 205 nm (ASL) (Kent et al. 1988).

Acid-insoluble Residue (AIR)

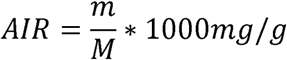

Where m= weight of the residue after drying

M= Oven dry weight of the sample before acid hydrolysis.

Acid-Soluble Lignin (ASL)

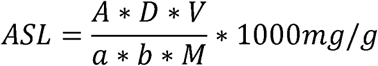

Where A=Absorbance,

D=Dilution factor,

V=Volume of the filtrate,

a=Extinction co-efficient of lignin,

b=Cuvette path length and

M=Oven dry weight of the sample before acid addition.

Total Lignin Content = AIR+ASL.

## Preparation of extracts for anti-microbial activity

*Coleus* leaves, root and stem samples were washed thoroughly under running tap water and then with distilled water to remove the dirt and to reduce the microbial load. The plant materials were air-dried under shade away from sunlight for 4-5 days, made into a fine powder using mortar and pestle. Extracts were prepared using polar solvent ethanol and non-polar solvent chloroform at a concentration of 10 g in 100 mL of solvent, allowed for the extraction of secondary metabolites with vigorous shaking for 48-72 h. The extracts were filtered and concentrated using Rota-evaporator which can be further diluted to the required concentration in DMSO used for assessing their anti-microbial activities by studying minimum inhibitory concentration (MIC) against bacterial strains *Escherichia coli* (MTCC 1652)*, Staphylococcus aureus* (MTCC 3160)*, Pseudomonas aeruginosa* (MTCC 1688)*, Bacillus cereus* (MTCC 430) and fungal strains *Aspergillus niger* (MTCC 282)*, Aspergillus flaws* (MTCC 873)*, Fusarium oxysporum* (MTCC 6659) and *Rhizopus stolonifer* (MTCC 2591) obtained from Microbial Type Culture Collection Centre, Institute of Microbial Technology (IMTECH), Chandigarh, India.

## Preparation of Inoculum

The colonies of test organisms grown overnight were inoculated into 0.85% normal saline and the turbidity adjusted to 0.5 Mc Farland using the standard which is equal to 1.5×10^8^ CFU/ml. It was further diluted to obtain the final inoculum of 5× 10^5^ CFU/ml.

## Determination of antimicrobial activity by minimum inhibitory concentration (MIC) method

MIC was performed as per Clinical and Laboratory Standards Institute guidelines using *Coleus* extracts against bacterial and fungal pathogens in a 96 well u-bottomed microtitre plates using p-iodonitrotetrazolium violet as an indicator dye. The ethanol and the chloroform extracts of *Coleus* was serially diluted from the concentration of 500 mg/ml to 0.02 mg/ml and then added with the final inoculum of 5× 10^5^ CFU/ml. The anti-microbial compound and the final inoculum were in the ratio of 1:1 (v/v). Each test performed in triplicate with positive and negative controls. After the addition of inoculum, plates were sealed with aluminium foil and incubated at 37°C for 24 h in the case of bacterial cultures and for 48 h at 28°C for fungal cultures respectively in an incubator. At the end of incubation period, the wells were added with 40 µL of 0.2 mg/ml p-iodonitrotetrazolium violet dye and incubated for 30 minutes for the colour development. Presence of bacterial or fungal growth is indicated by a change in the colour of the medium to red, whereas no colour change indicates the absence of growth of the organism and the least concentration where there is no growth is considered as an MIC value of that particular compound against bacterial and fungal strains used. Ampicillin and Fluconazole were used as standards.

## Statistical analysis

Results mentioned are reported as the mean ± standard error (SE) values of five independent experiments, conducted on five different plants in each experiment. SE values were calculated directly from the data according to standard methods (Taylor 1982). Data analysis was carried out using the SPSS package. Mean values were compared by Duncan’s multiple range test and *P*-values which are less than or equal to 0.05 were considered as statistically significant.

## Results

Quantitative determination of ten different secondary metabolites namely phenols, flavonoids, tannins, lignins, alkaloids, steroids, cardiac glycosides, anthraquinones, terpenes and saponins were carried out in leaf, stem and root samples of *Coleus* species (Fig. 1-10). The range of secondary metabolites in leaf, stem and root of *Coleus* species was found to be 0.75-3.82 mg/g for phenols, 0.3-0.95 mg/g for flavonoids, 0.88-2.62 mg/g for tannins, 0.11-2.4 mg/g for cardiac glycosides, 0.14-0.94 mg/g for anthraquinones, 11.4-31% for lignins, 3.91-6.2 mg/g for steroids, 0.9-3.82 mg/g for saponins, 2.3-9.2 mg/g for alkaloids and 123-315 mg/g for terpenes. The concentration of bioactive compounds varies among the species and within the species under saline conditions. The amount of plant bioactive compounds increased with the increasing concentration of NaCl up to the optimum level and the amount decreased with the increasing concentrations of NaCl beyond the optimum level. In the present study, the content of secondary metabolites in *Coleus* has increased under mild (100 mM), moderate (200 mM) and severe (300 mM) salinity treatment (Fig. 1-10). Thereafter, decrease in the level of secondary metabolites at the concentration above 300 mM NaCl is noticed and the experiment was designed considering the salinity treatment up to a concentration of 300 mM NaCl. The maximum increase in the level of bio-active compounds was observed at a concentration of 300 mM NaCl. The content of terpenes was found to be higher in all the five *Coleus* species compared to other bioactive compounds. The concentration of the majority of the secondary metabolites were found to be high in leaf samples of *Coleus* followed by stem and root, whereas few bioactive compounds were high in stem compared to leaf and root of *Coleus* species.

**Fig. 1.**
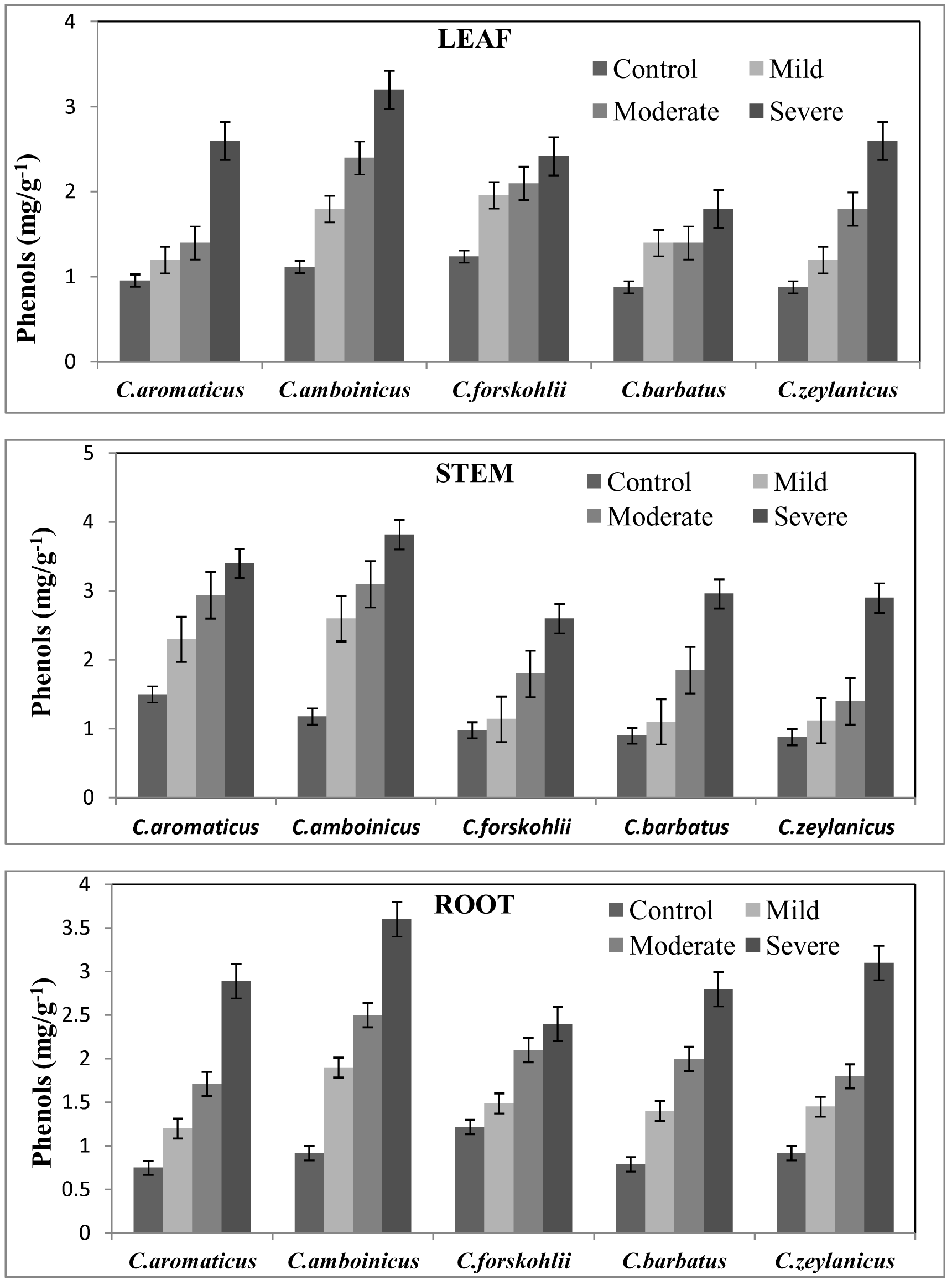
Quantitative determination of phenols in leaf, stem and root of five different *Coleus* species under normal and saline conditions. Each point is an average of five independent determinations ± SE, (t _(4)_ = 0.1, *p* □ 0.05).

**Fig. 2.**
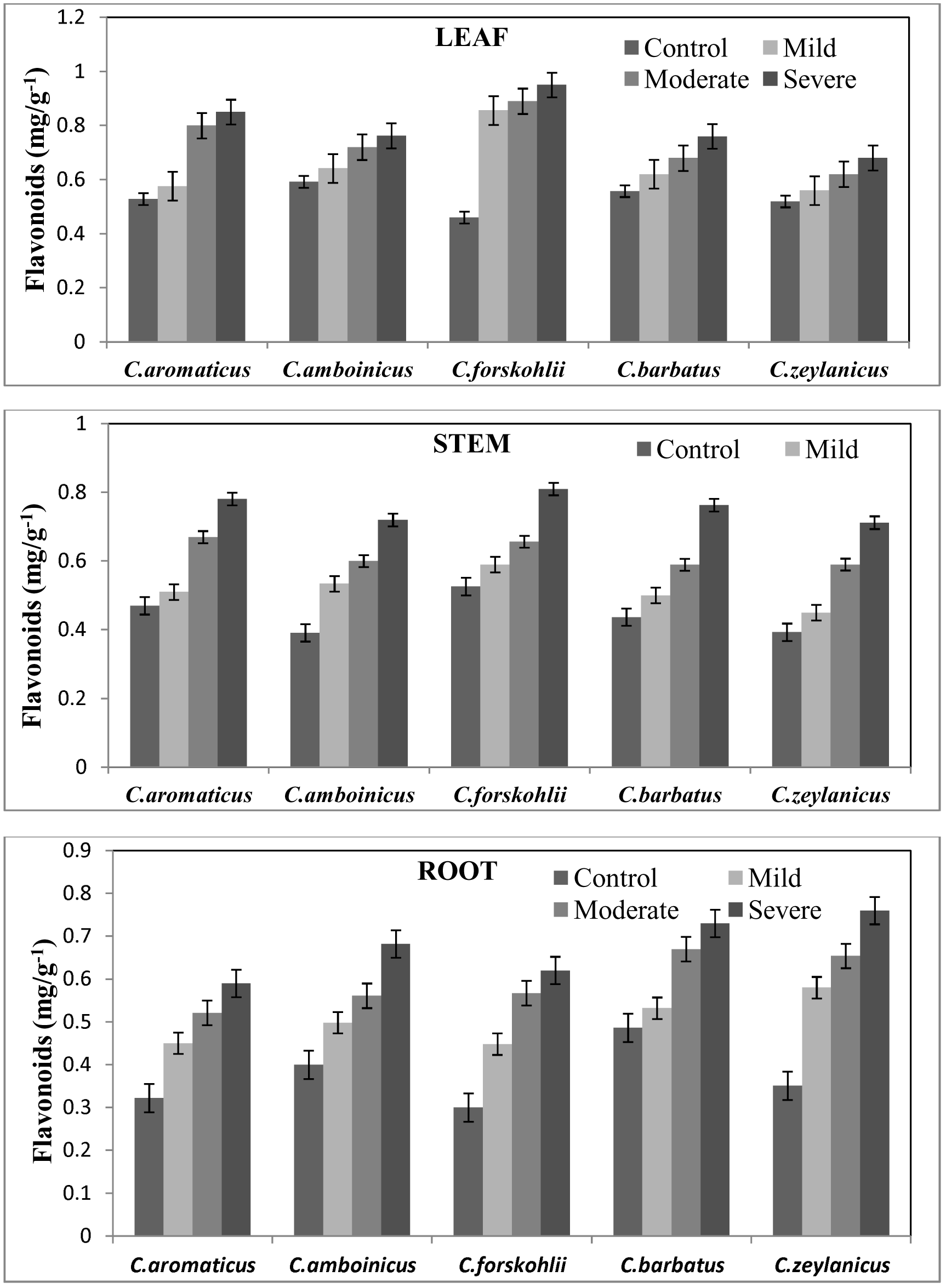
Quantitative determination of flavonoids in leaf, stem and root of five different *Coleus* species under normal and saline conditions. Each point is an average of five independent determinations ± SE, (t _(4)_ =0.10, *p □* 0.05).

**Fig. 3.**
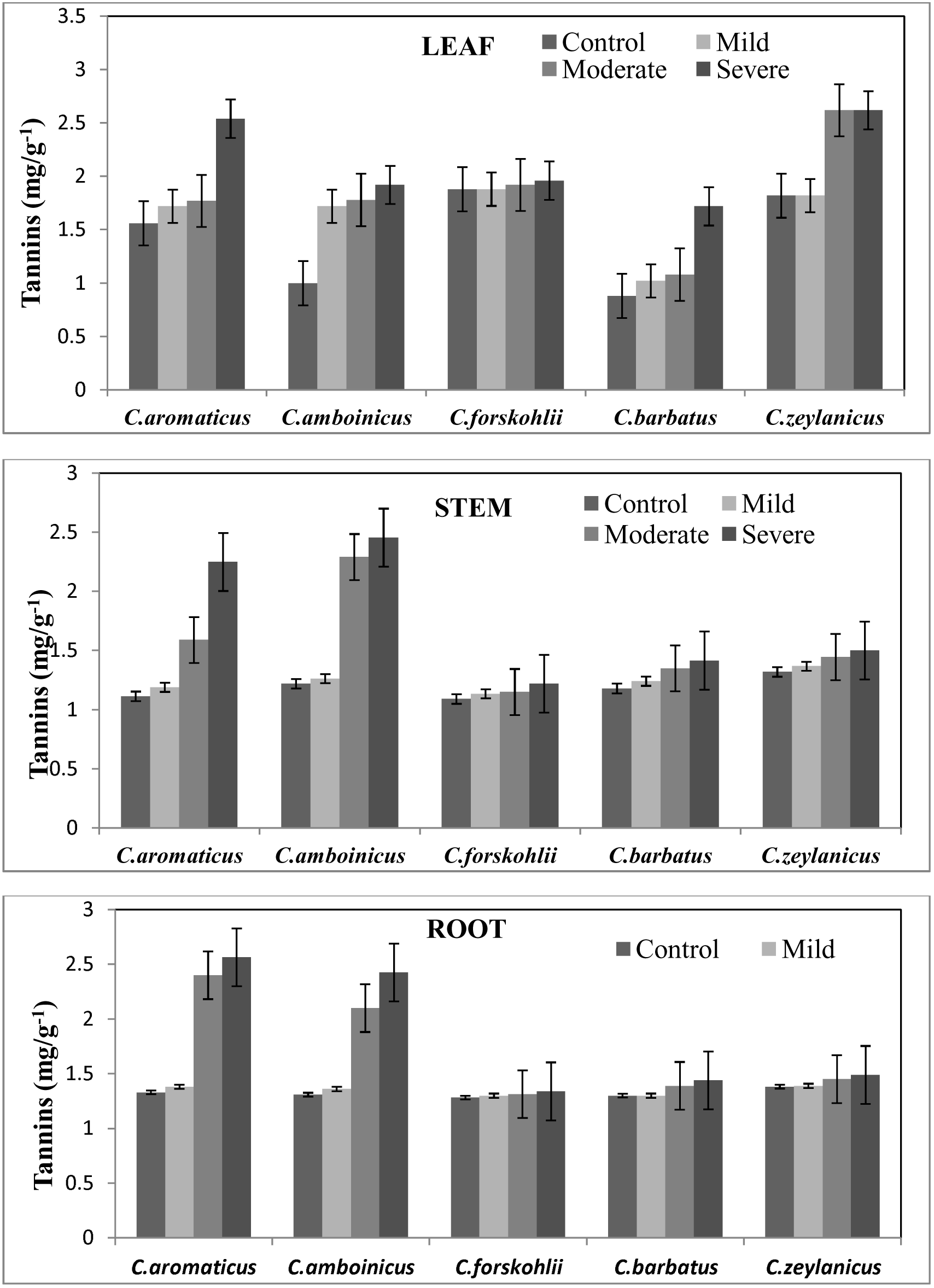
Quantitative determination of tannins in leaf, stem and root of five different *Coleus* species under normal and saline conditions. Each point is an average of five independent determinations ± SE, (t _(4)_ = 0.28, *p □* 0.05).

**Fig. 4.**
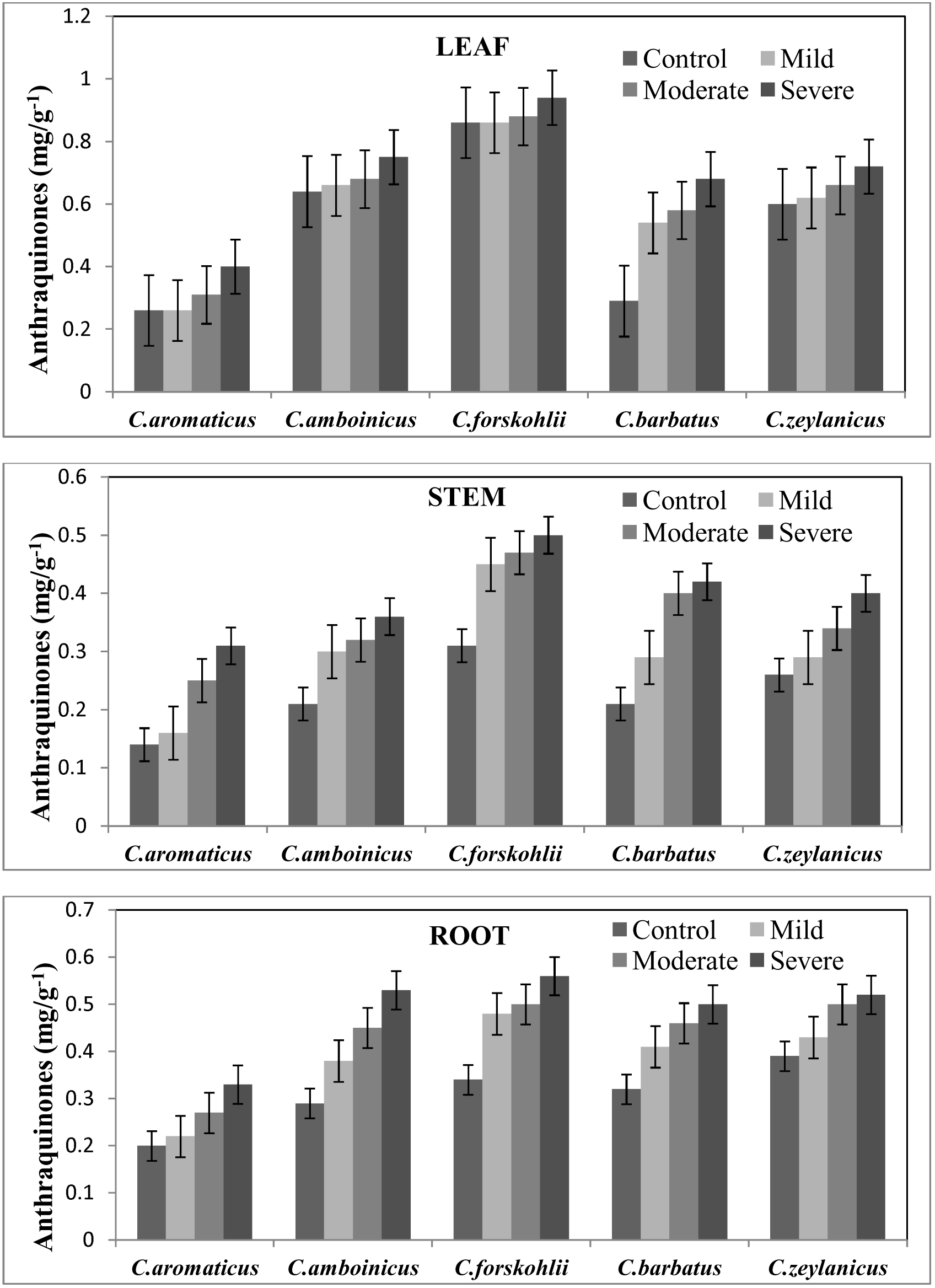
Quantitative determination of anthraquinones in leaf, stem and root of five different *Coleus* species under normal and saline conditions. Each point is an average of five independent determinations ± SE, (t _(4)_ =0.32, *p □* 0.05).

**Fig. 5.**
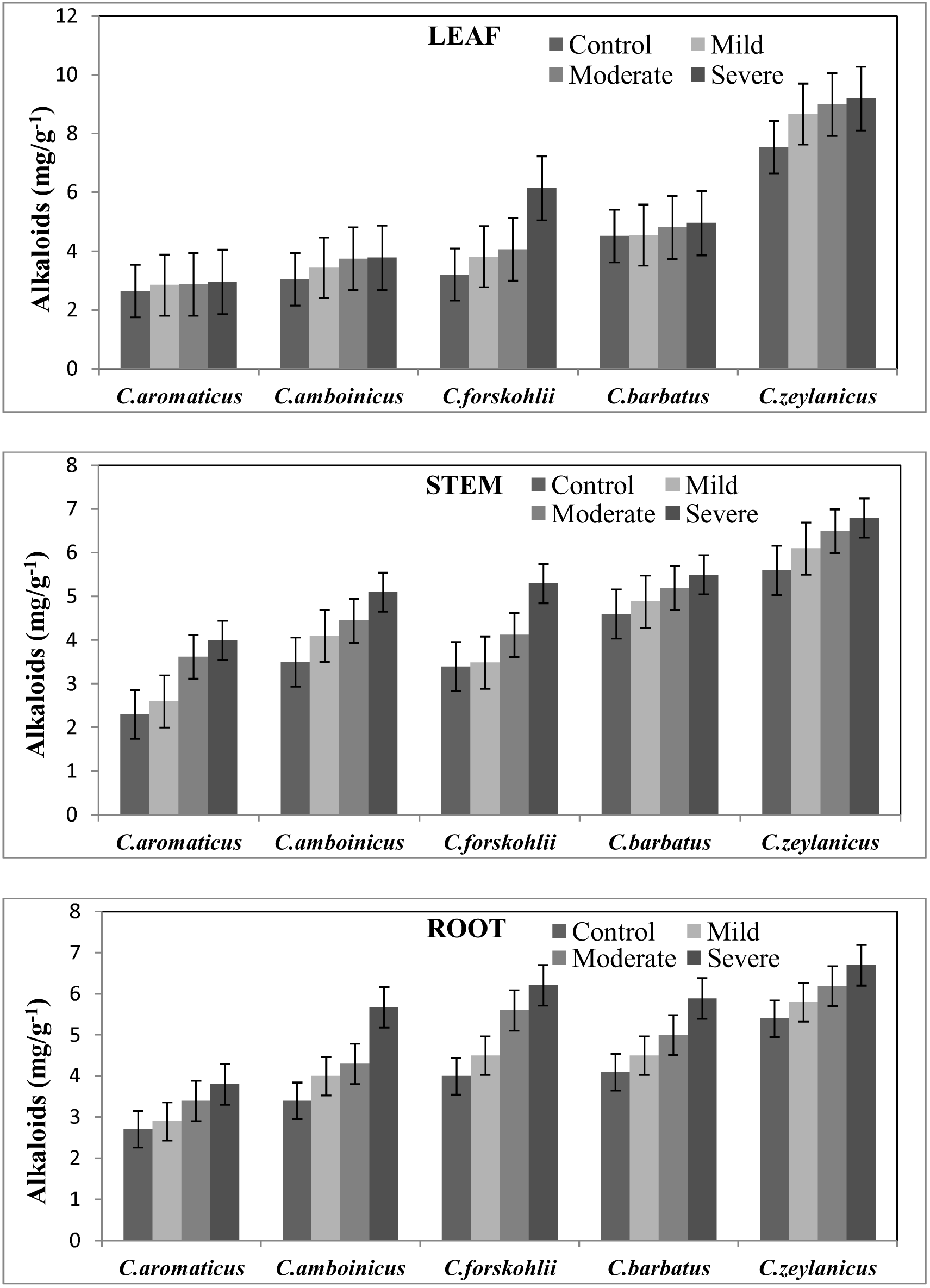
Quantitative determination of alkaloids in leaf, stem and root of five different *Coleus* species under normal and saline conditions. Each point is an average of five independent determinations ± SE, (t _(4)_ = 0.46, *p □* 0.05).

**Fig. 6.**
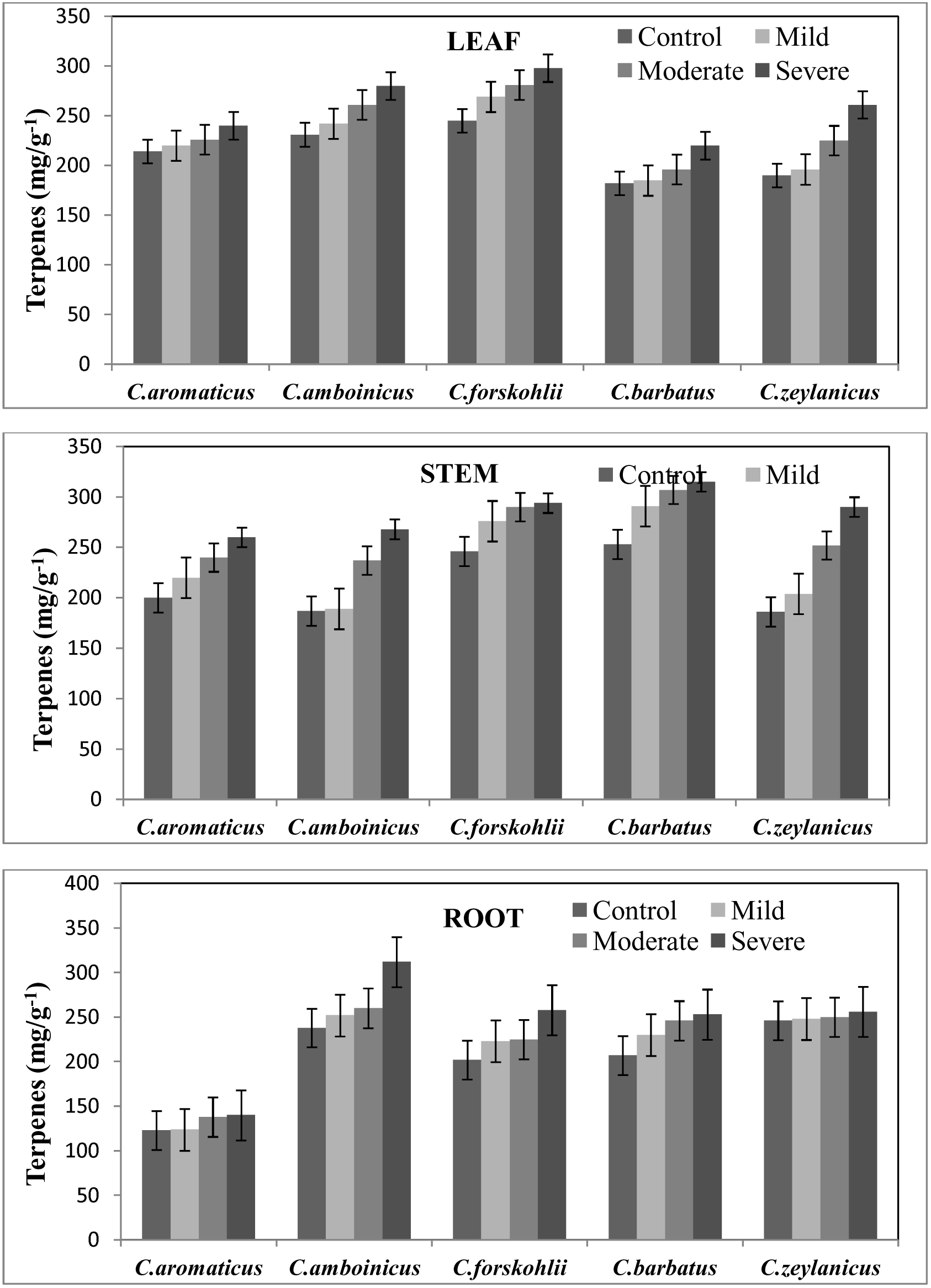
Quantitative determination of terpenes in leaf, stem and root of five different *Coleus* species under normal and saline conditions. Each point is an average of five independent determinations ± SE, (t (_4_) = 3.14, *p* □ 0.05).

**Fig. 7.**
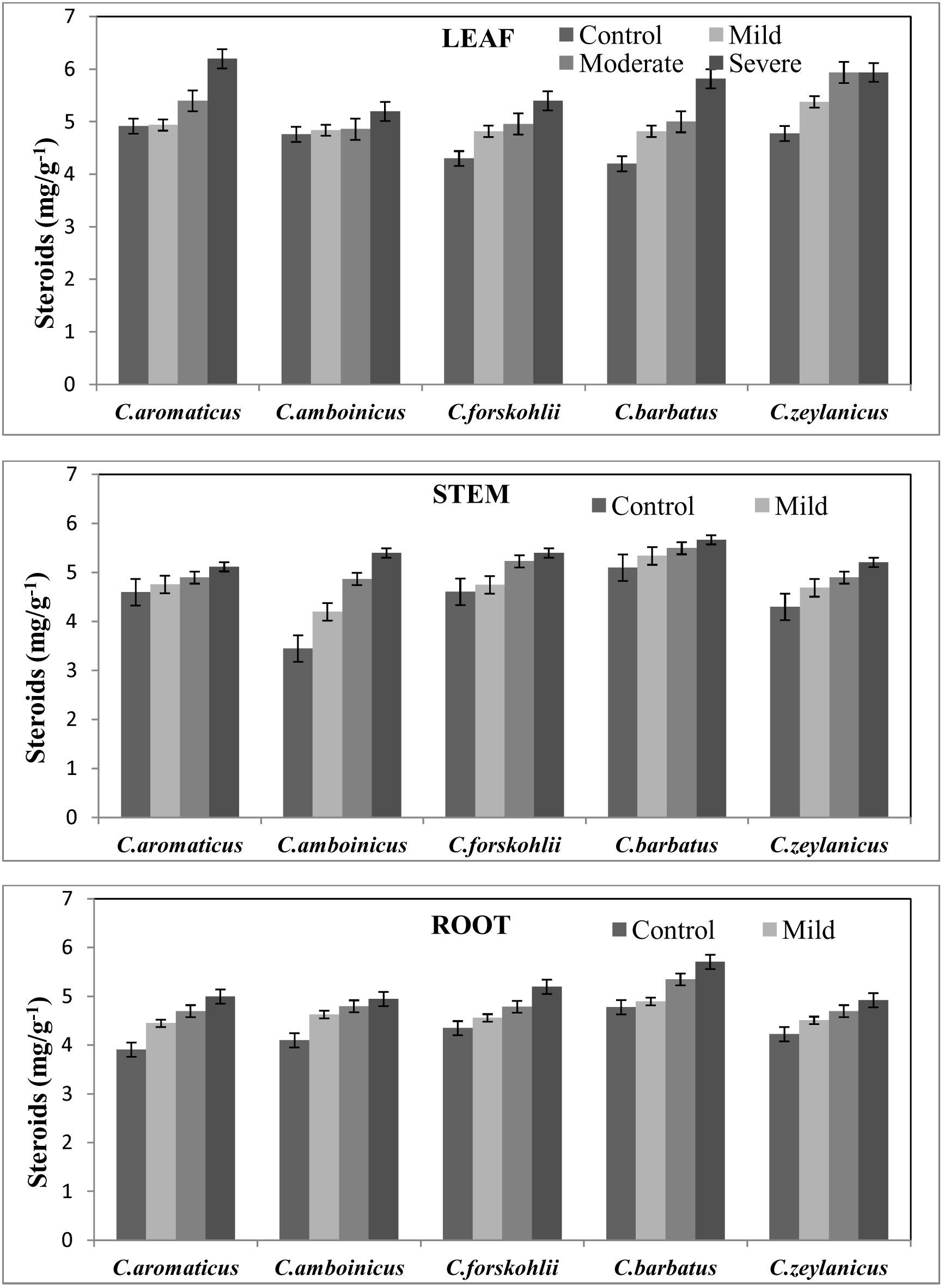
Quantitative determination of steroids in leaf, stem and root of five different *Coleus* species under normal and saline conditions. Each point is an average of five independent determinations ± SE, (t _(4)_ = 0.92, *p □* 0.05).

**Fig. 8.**
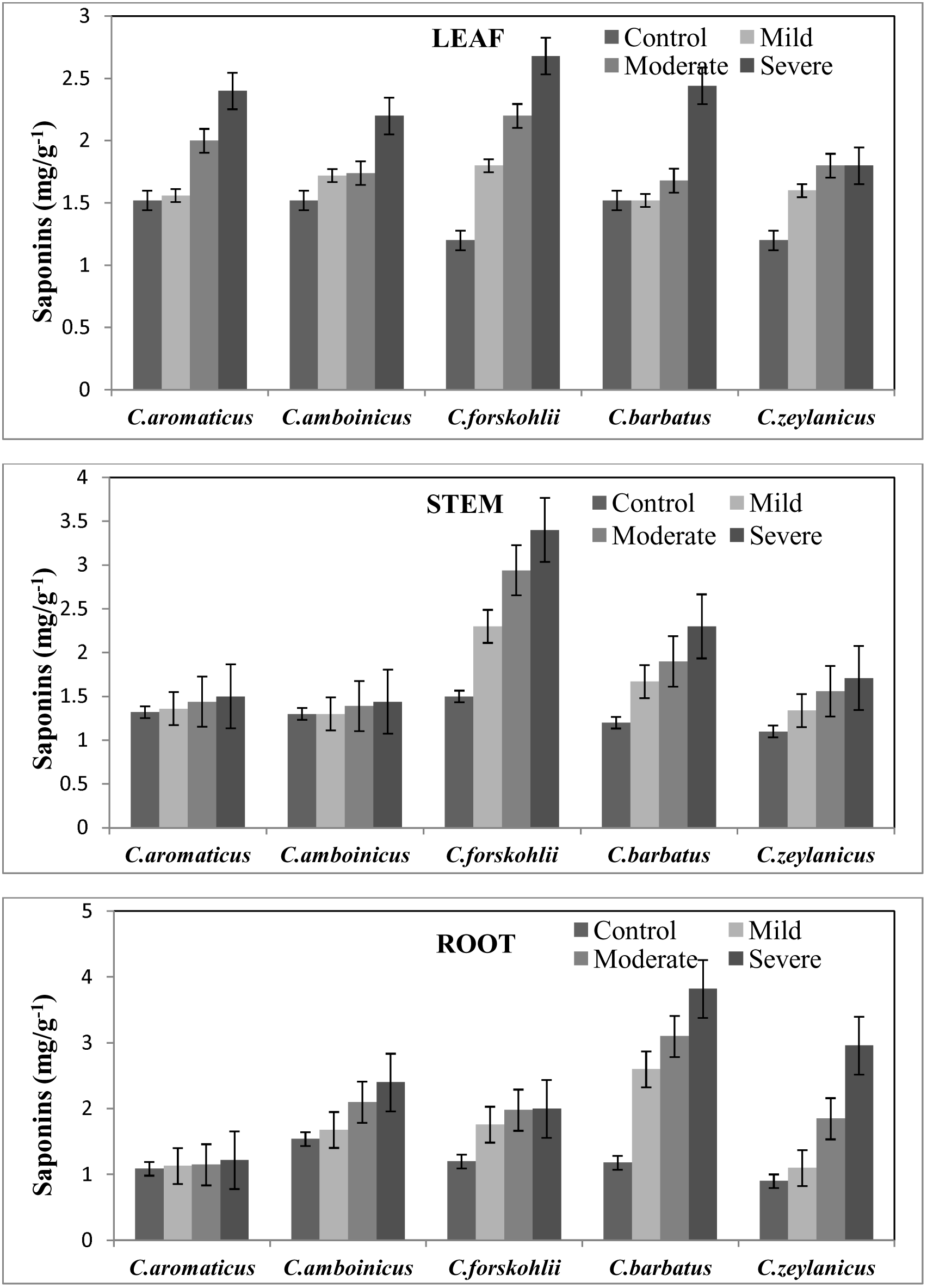
Quantitative determination of saponins in leaf, stem and root of five different *Coleus* species under normal and saline conditions. Each point is an average of five independent determinations ± SE, (t _(4)_ =0.4, *p □* 0.05).

**Fig. 9.**
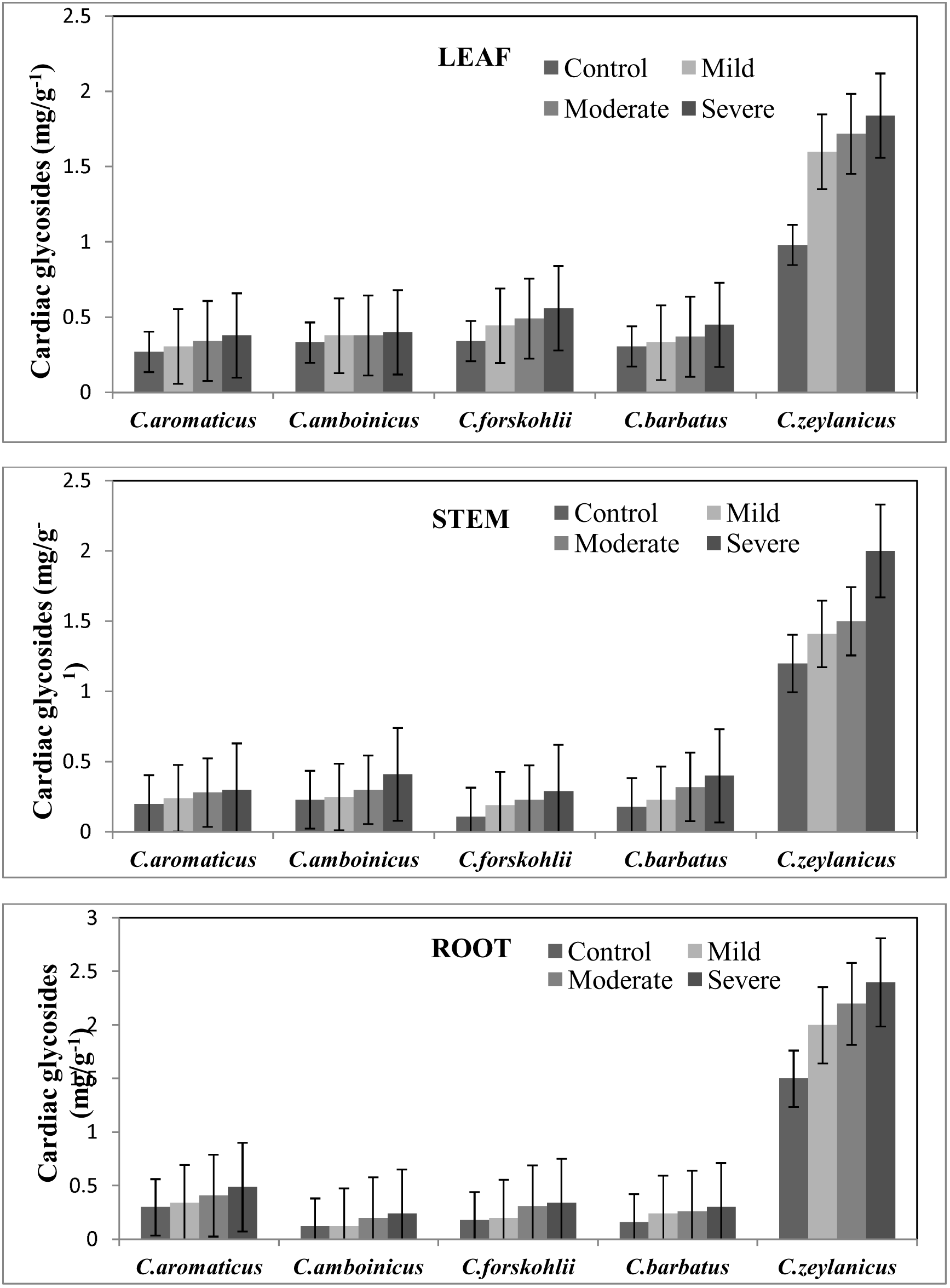
Quantitative determination of cardiac glycosides in leaf, stem and root of five different *Coleus* species under normal and saline conditions. Each point is an average of five independent determinations ± SE, (t _(4)_ =0.1, *p* □ 0.05).

**Fig. 10.**
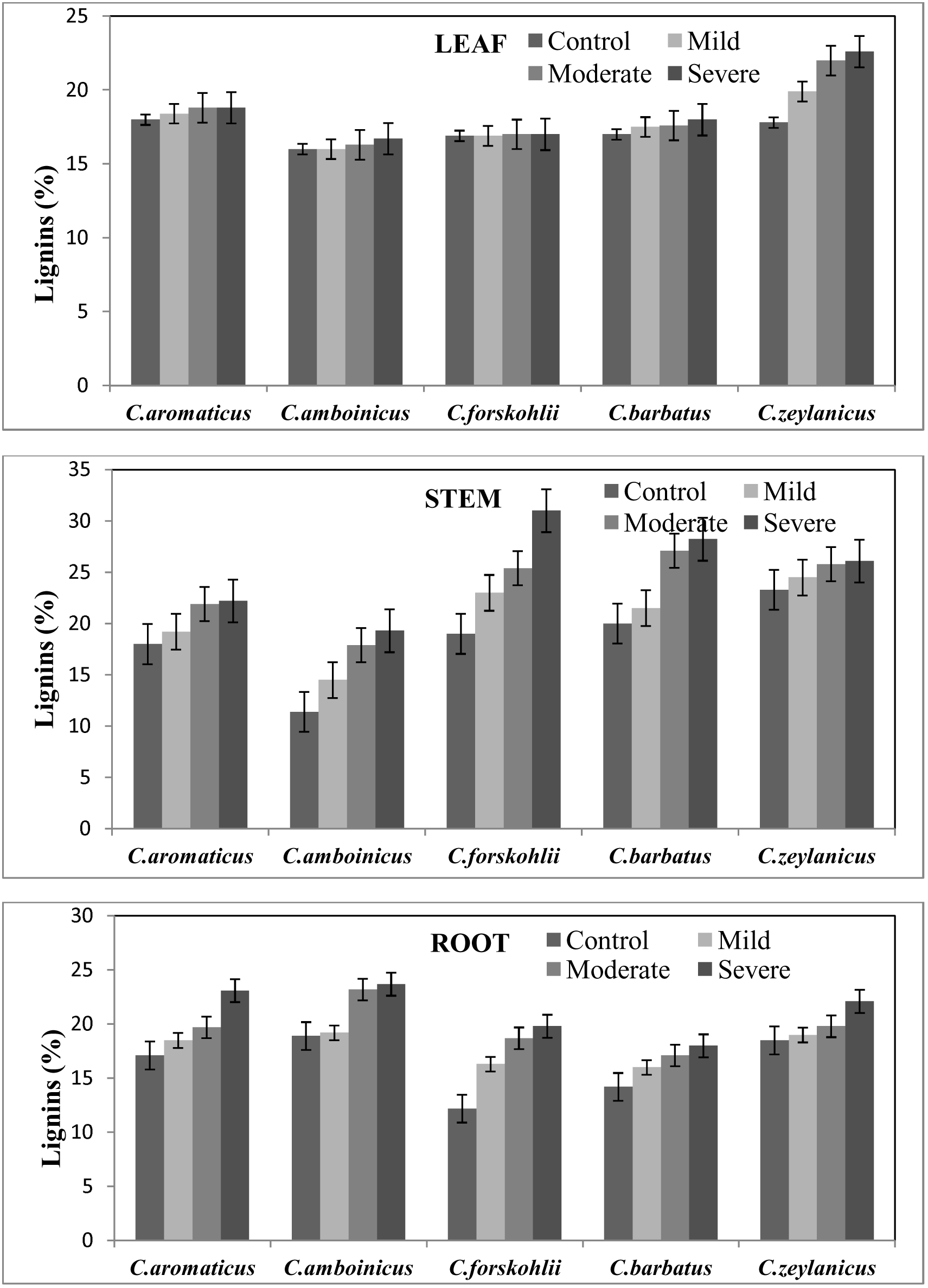
Quantitative determination of lignins in leaf, stem and root of five different *Coleus* species under normal and saline conditions. Each point is an average of five independent determinations ± SE, (t _(4)_ = 0.80, *p □* 0.05).

The effect of salt stress on anti-microbial activity of five different *Coleus* species, namely *C.aromaticus, C.barbatus, C.amboinicus, C.forskohlii* and *C.zeylanicus* ethanol and chloroform extracts against four bacterial strains *Escherichia coli, Bacillus cereus, Staphylococcus aureus, Pseudomonas aeruginosa,* and four fungal strains *Aspergillus niger, Aspergillus flavus, Rhizopus stolonifer* and *Fusarium oxysporum* is depicted in Table 1-2. The leaf, stem and root extracts of all the five *Coleus* species showed good anti-microbial activity against tested pathogenic strains by inhibiting their growth. The leaf extracts of *Coleus* showed higher inhibitory activity against tested strains followed by the stem and root extracts. Ethanol extracts showed high anti-microbial activity ranging from 1.5-100 mg/ml compared with the chloroform extracts ranging from 0.97-250 mg/ml against tested bacterial and fungal pathogens respectively and the activity increased with increasing salinity due to the up regulation of secondary metabolites whereas for few species of *Coleus* against few tested strains, the activity remained to be the same as control values. *Bacillus cereus* was highly susceptible bacterium whose activity was inhibited at a concentration of 1.5 mg/ml and 0.97 mg/ml by *Coleus forskohlii* ethanol and chloroform extracts whereas, *A.niger* was the highly susceptible fungus inhibited by *Coleus zeylanicus* and *Coleus forskohlii* leaf extracts at a concentration of 0.39 mg/ml respectively. Among the five different *Coleus* species used in the study, *C.forskohlii* showed high anti-microbial activity both under normal and saline conditions followed by *C.zeylanicus.*

**Table 1.**
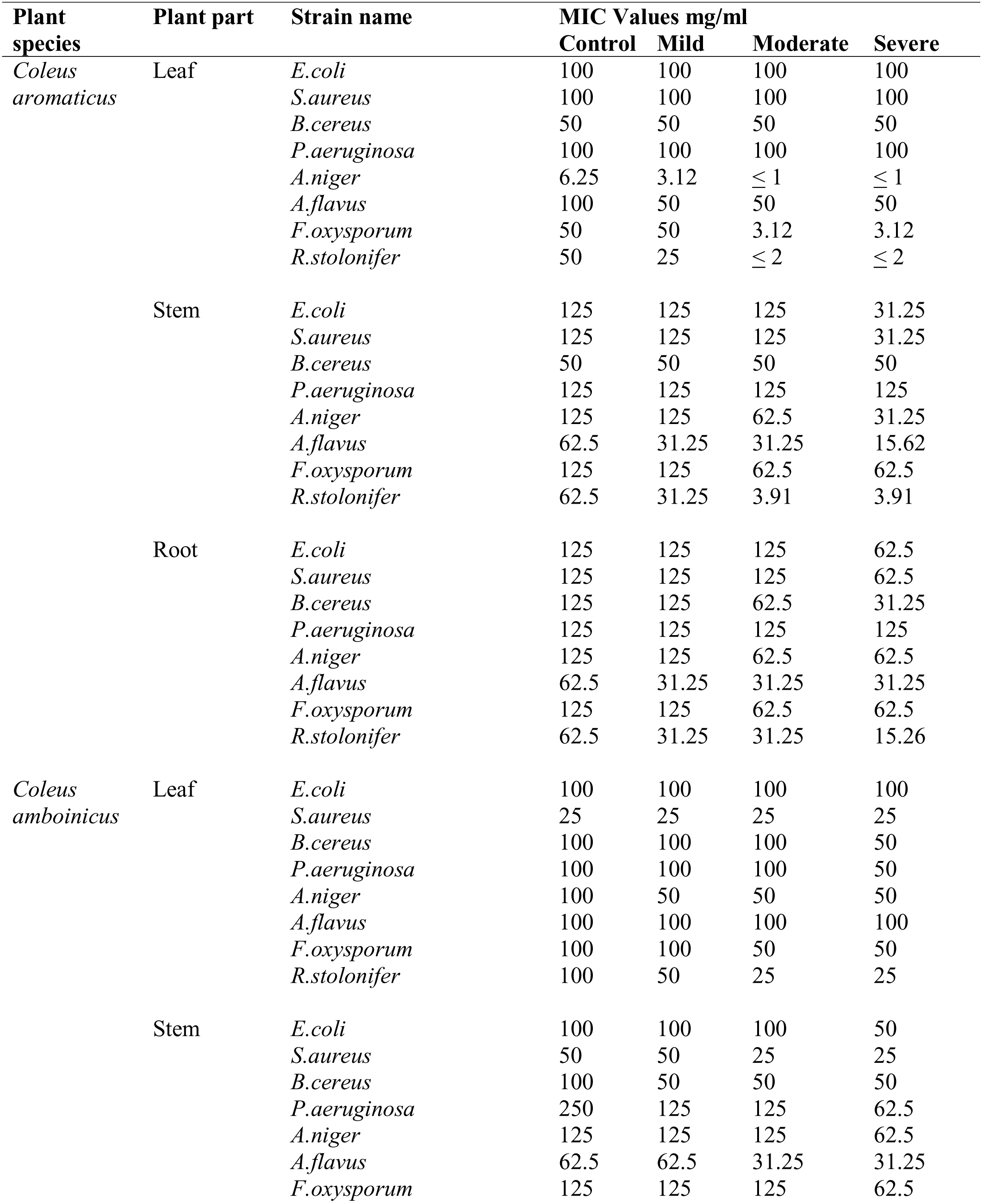
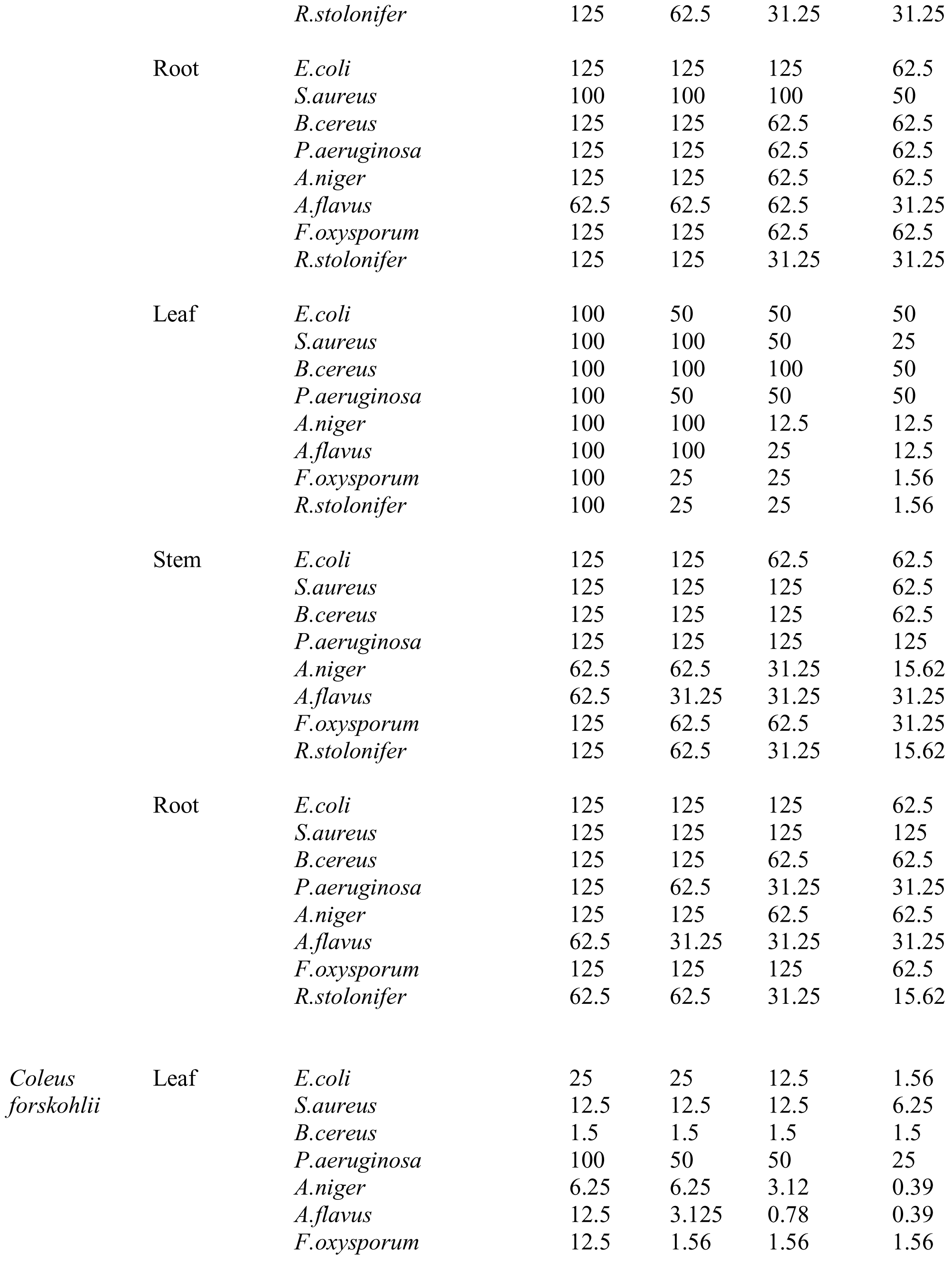
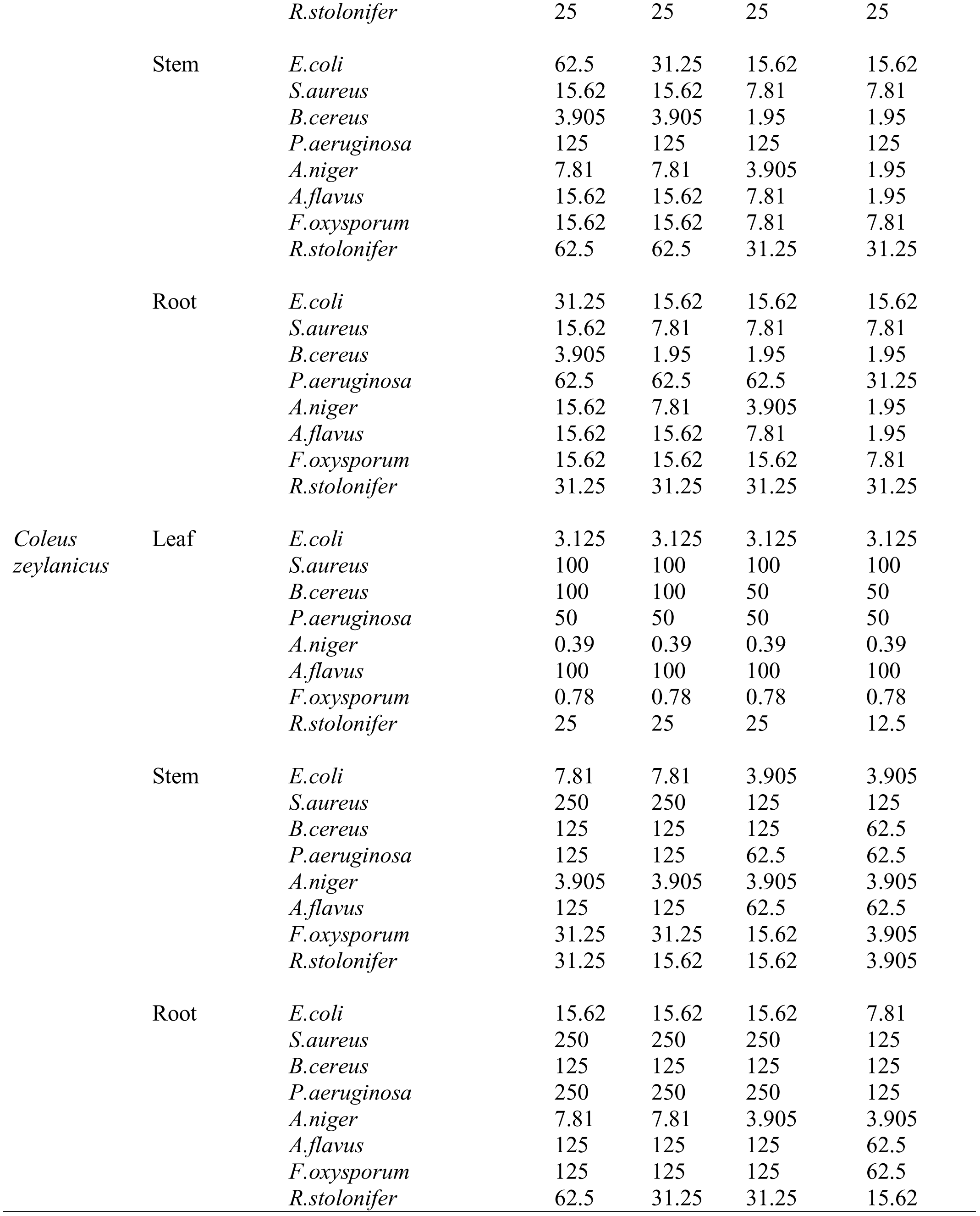
Anti-microbial activity of *Coleus* ethanol leaf, stem & root extracts in mg/ml.

**Table 2.**
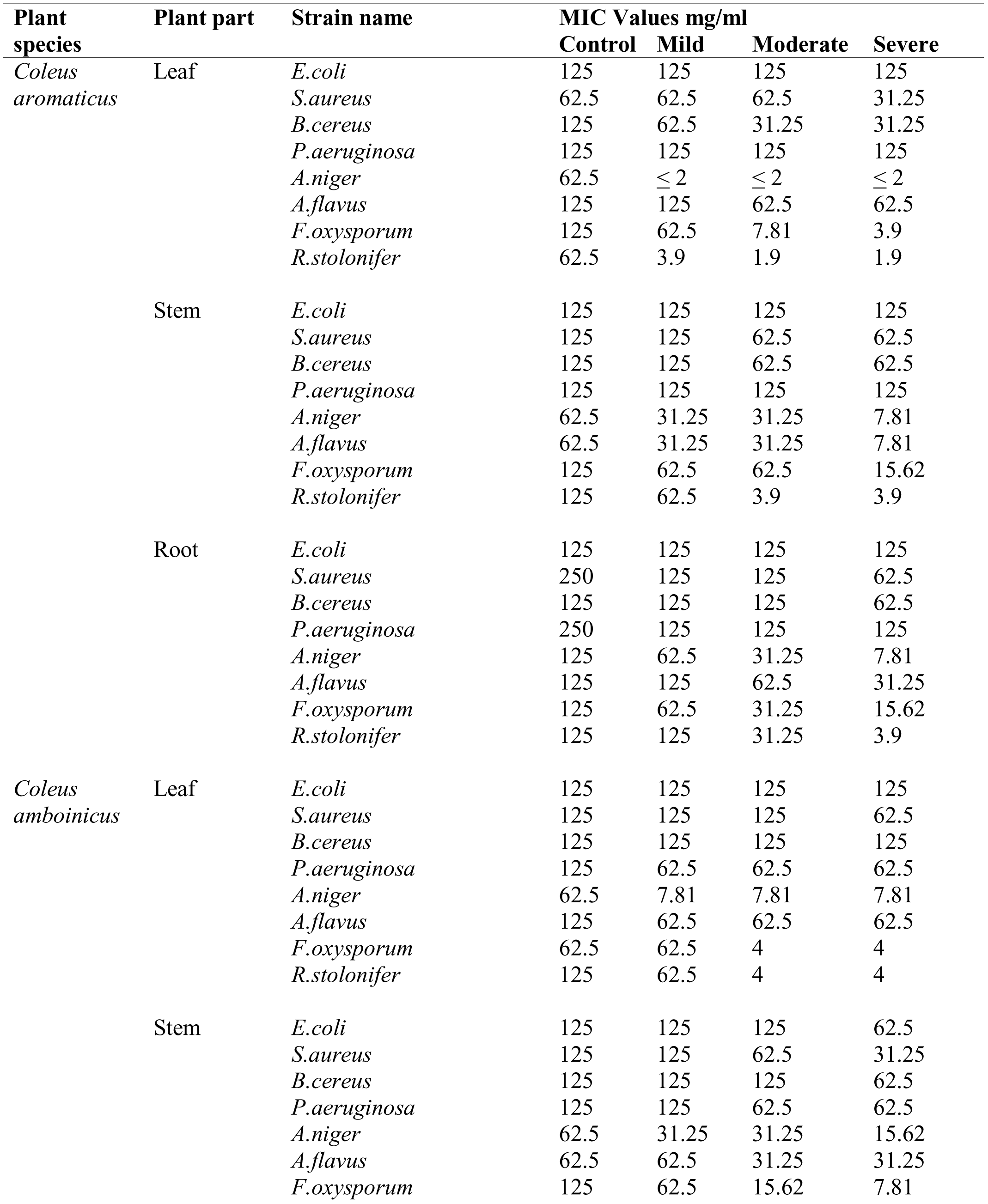
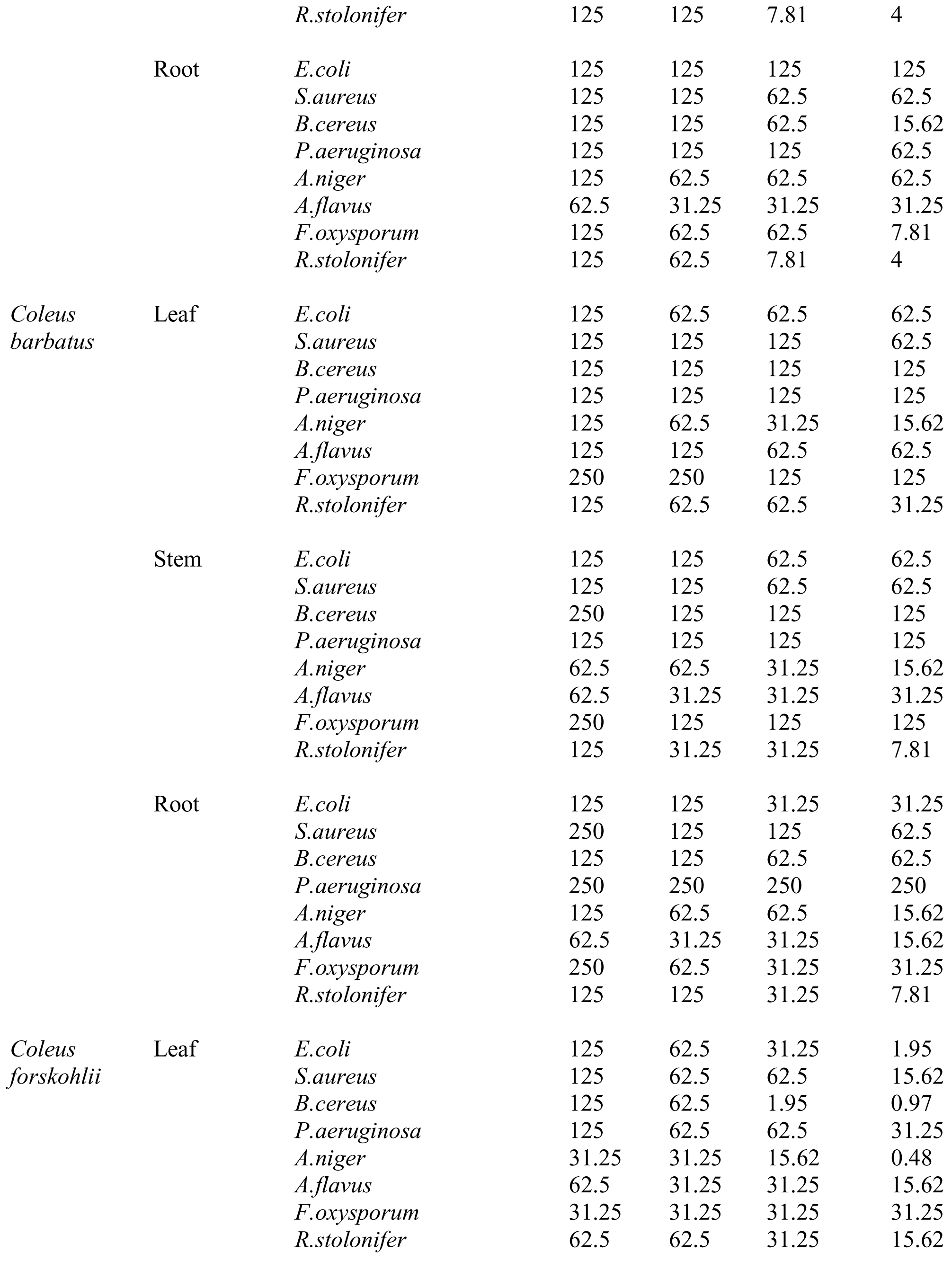
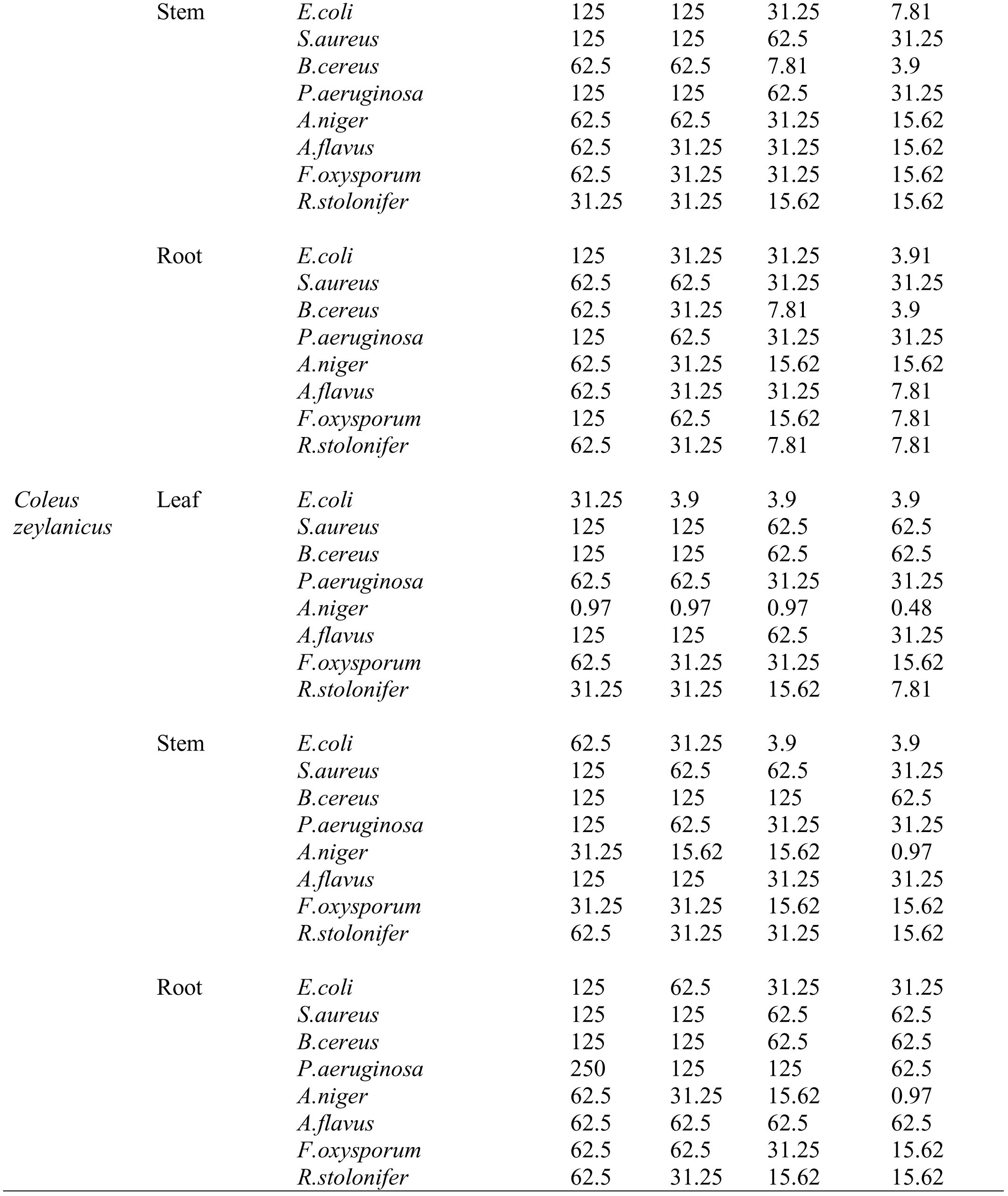
Anti-microbial activity of *Coleus* chloroform leaf, stem & root extracts in mg/ml.

## Discussion

Medicinal plants produce a large number of secondary metabolites with several biological properties. The presence of polyphenols and their up regulation during stress play a key role in the plant defense mechanisms. An extensive study on phytoconstituents has been made in five different species of *Coleus* leaf, stem and root tissues subjected to salinity stress. The increase in the content of phenolic compounds with increased salinity was observed during the study and our data is supported by the findings of Valifard et al. (2014); reported the increased total phenolic compounds in the leaf samples of medicinal, aromatic plant *Saliva mirzayanii* under salinity stress. The increased total phenol under moderate salinity stress in the red pepper plant was reported (Navarro et al. 2006). In our present study, the increase in the content of secondary metabolites is seen in leaf, stem and root samples of *Coleus* under salinity stress, whereas, the concentration of secondary metabolites were found to be high in leaf of *Coleus* followed by stem and root. The increase in the content of phenolic compounds with increased salinity in different parts of the plant was reported (Muthukumarasamy et al. 2000). The growth of the plant during salinity stress is reduced due to the accumulation of toxic ions, Na^+^ and Cl^-^ (Marosz and Nowak 2008). The increase in the vacuolar volume mediates directional expansion causes primary plant cell growth and also facilitates the osmotic adjustment essential for cellular development by compartmentalization of Na^+^ and Cl^-^ (Shuji et al. 2002). By decreasing the leaf area, plant tries to cope with the condition of abiotic stress thereby conserving the energy. The plant potassium nutrition is disrupted by the sodium ions at the surface of root because of the similarity between potassium and the sodium ions; the uptake of potassium by root system is strongly inhibited. The uptake of potassium by plants take place either by the high affinity or low affinity system, but generally plants undergo high affinity potassium uptake system during salinity stress to maintain appropriate potassium nutrition to maintain enzyme activities, cell turgor and membrane potential as the deficiency leads to the reduced plant growth.

Plants when exposed to abiotic stress like salinity stress, their growth will be reduced and generate a high oversupply of reduction equivalents. The massive amounts of NADPH^+^, H^+^ (strong reduction power) enhance the synthesis of compounds like alkaloids or phenols and isoprenoids which are highly reduced. The massive generation of oxygen radicals and the damage by photo-inhibition is prevented by the accumulated secondary metabolites or natural products of plants affected by stress (Xin et al. 2011). The enhanced levels of secondary metabolites during salinity stress might be due to the inductions in enzymatic activity favouring the production of different bioactive compounds. The presence of alkaloids in *Coleus* might be responsible for anti-malarial, analgesic activity and its use in the treatment of stomach disorders. Similar results of higher alkaloid content in the salinity treated plants of *C.roseus* compared to control plants were reported (Abdul et al. 2008). Tannins used to heal inflamed mucous membrane and wounds due to its astringent property. Bioactive compounds like terpenes, steroids and saponins possess cardiac and hypertensive depressant activity. Terpenoids possess anti-cancer properties, promotes apoptosis. The concentration of terpenoids was found to be high in *Coleus* plant which might be the reason of anti-cancer potential. Similar range of 216.67 to 350 mg/g terpenoids were reported in *Ocimum* (Vimala et al. 2014). The presence of cardiac glycosides found to be effective in congestive heart failure (Aboaba et al. 2001). Flavonoids are one of bioactive compounds accumulate and trigger the synthesis of substances with defensive role. The anti-viral, anti-inflammatory and antioxidative properties of medicinal plants are due to the presence of flavonoids, which are used to treat several conditions like diabetes, ulcers, rheumatic fever and hypertension. Kidney disorders and stomach problems can be cured with the use of plant polyphenols (Vimala and Francis 2015). The presence and the composition of different bioactive compounds in medicinal plants is controlled both at the environmental and genetic level (Awika and Rooney 2004). The demand for the use of medicinal plants rich in phenolics in food industries is increasing because of their ability to improve the quality and nutritional value of foods. These compounds contain hydroxyl groups which can degrade lipids and scavenge free radicals (Naima Saeed et al. 2012). The laxative property of anthraquinones is generally used in pulp bleaching for production of paper as it is a building block for most of the dyes (Soladoye and Chukwuma 2012). The important role of conducting water in the stem of plants is done by lignins. From our study, it was observed that all the five *Coleus* species were tolerant to salinity stress, acquiring resistance to salinity by the accumulation of secondary metabolites thereby providing the osmotic balance to the plant and by protecting the cells, preventing the damage caused by the generation of oxygen radicals.

In recent years, a number of multi-drug resistant strains have developed by expressing resistant genes due to the improper usage of antibiotics. To avoid this problem, there is a need to develop new alternate drugs to eradicate the pathogenic population. Medicinal plant extracts with antimicrobial activity can be used as a desirable tool to eradicate the population of pathogenic strains, particularly in the treatment of infectious, dreadful diseases and in food spoilage. The initial step for the discovery of new drugs with antimicrobial potential is the screening of plant extracts (Cseke et al. 2016). Among the different parts of the plant, leaf is considered to be one of the highest accumulator regions for compounds used generally for therapeutic needs (Jagtap et al. 2009). In the present study, control, mild, moderate and severe Nacl treated species of *Coleus* leaf, root and stem ethanol and chloroform extracts were tested against four bacterial strains and four fungal strains, which have inhibited their growth. Antimicrobial activities of *Coleus* extracts might be due the presence of various bioactive compounds exhibiting antiviral, antimicrobial, anti-inflammatory and antioxidant properties. *Coleus* leaf, stem and root extracts have shown effective antimicrobial activity against gram positive, gram negative and fungal strains used in the study which indicates the presence of antimicrobial compounds exhibiting broad spectrum activity. The activity of microbial growth inhibition increased with increased salinity is due to the up regulation of plant bioactive compounds. The difference in the antimicrobial activity of leaf, stem and root extracts of *Coleus* is due to the difference in the composition and the concentration of phytochemicals present within a particular tissue. The effect of salt stress and the type of solvent used for the extraction also influence the antimicrobial activity. The antimicrobial activity varies with the species to species or within the species is due to variations in the secondary metabolite profiles and various other factors like climatic and environmental changes. The response of plants to produce one metabolite over the other is due to the effect of different stress factors. The composition of plant secondary metabolites is altered due to the difference in the level of carbon dioxide, altitude and the presence of pathogenic microbes and insects (William et al. 2016). The inhibitory effect of *Coleus* leaf, root and stem ethanol and chloroform extracts on the tested bacterial strains ranged from 250-1.5 mg/ml, whereas, against fungal strains MIC values ranged from 150-0.39 mg/ml respectively. Similar results of antimicrobial activity of *Coleus barbatus* ethanol and chloroform extracts against the strains of *S.aureus* and *P.aeruginosa* were reported by Abhishek et al. (2011). The inhibitory activity at a concentration of 100 mg/ml against the strains of *E.coli* and *S.aureus* by *Moringa oleifera* leaf ethanol extract was reported by Ibrahim et al. (2015). Jacqueline et al. (2017) reported the antifungal activity of *Coleus* species methanol extracts against the strains of *Aspergillus, Rhizopus, Mucor, Rodotorula, Geotricum, Brasidiobolus, Trichophyton, Microsporum, Epidermophyton* and *Candida* support our study which states the antifungal potential of *Coleus* extracts. The presence of secondary metabolites in *Coleus* species plays a major role in protecting the plant from stress also responsible for the anti-microbial activity. It was believed that the extracts exhibit antimicrobial potential of causing damage to the nucleotides with increased spatial division and by genetic material condensation (Thilagavathi et al. 2016). The action of bioactive compounds on microbes might be due to the interference of bacterial cell wall peptidoglycan biosynthesis and by inhibiting protein synthesis, nucleic acid synthesis, act as chelating agents, inhibiting the metabolic pathway, disrupting the peptide bonds and preventing the microbes to utilize the available nutrients. The leaf extracts of *Coleus* showed potent inhibitory activity compared to stem and root might be due to the presence of number of bioactive compounds with antimicrobial property. The secondary metabolites are generally deposited in different parts of the plant in different proportions of an individual plant as the production of phytochemicals in leaves is expected to be higher compared to the other parts of the plant (Clarice et al. 2017). The growth and the metabolism of microorganisms are inhibited by the interference of the active components present within a bio-active compound (Aboaba et al. 2006). *Bacillus cereus* was found to be the most susceptible bacterium inhibited at a concentration of 0.97 mg/ml by *Coleus forskohlii.* Similar results of inhibitory activity on *Bacillus cereus* were reported by Abdelaaty et al. (2017). The difference in the antimicrobial activity between gram positive and gram negative strains is due to the difference in the composition of the cell wall. The extracts penetrate through the mesh like peptidoglycan layer of gram-positive microorganisms, whereas the penetration of extracts in gram negative strains is difficult as they possess outer lipopolysaccharide membrane. *Coleus* extracts effectively inhibited gram negative strains responsible for several infectious diseases in humans, therefore *Coleus* plant is considered to have high therapeutic value can be used in developing novel antimicrobial drugs to overcome the usage of conventional antibiotics. Many researchers have reported the broad spectrum antimicrobial activity of flavonoids, alkaloids, polyphenols and tannins. The tannins act by forming complex with polysaccharides, inactivating the enzymes, preventing microbial adhesion and precipitating the proteins (Prashant et al. 2017). From the above results, the whole *Coleus* plant is a good source of terpenoids, flavonoids and other secondary metabolites suggests the use of this herb in food and pharmaceutical industries. It was clear that the *Coleus* extracts possess metabolites effective in killing pathogenic microbes which can be used in the preparation of traditional medicine for therapy against several diseases.

## Conclusions

From the above results, it was clear that all the five *Coleus* species are capable of surviving during salinity stress up to the optimum levels of NaCl treatment with specific time period and with the up regulation of secondary metabolites possessing nutraceutical and pharmaceutical value for the development of new anti-microbial drugs against multi-drug resistant pathogenic strains to address unmet therapeutic needs. In addition, under salinity stress, an increase in the content of different bioactive compounds appears to be involved in the response of *Coleus* to NaCl stress and their presence responsible for the anti-microbial, anti-oxidant and anti-inflammatory properties of this medicinal plant. Thus, this medicinal plant can be considered in the development of new alternative traditional drugs in order to cure most dreadful diseases caused by the multi-drug resistant strains.

## Acknowledgements

Research lab of K.V. Chaitanya is funded by the grants from the University grants commission (UGC), Govt. of India, 42-197/2013. Divya is thankful for the UGC research fellowship.

## References

Abdelaaty A, Shahat, Elsayed A, Mahmoud, Abdullah A, Al-Mishari, Mansour S, Alsaid. 2017. Antimicrobial activities of some Saudi Arabian Herbal plants. African Journal of Traditional Complementary and Alternative Medicine. 14(2), 161–165.

Abdul Jaleel C, Beemarao S, Ramalingam S, Rajaram P. 2008. Soil salinity alters growth, chlorophyll content, and secondary metabolite accumulation in Catharanthus roseus. Turkish Journal of Biology. 32, 79–83.

Abhishek M, Rakshanda B, Prasad GBKS, Dua VK. 2011. Coleus barbatus as a Potent Antimicrobial Agent against Some Gastro-Intestinal Pathogens. Journal of Life Sciences. 3(2), 137–140.

Aboaba OO, Smith SI, Olide FO. 2006. Antimicrobial Effect of Edible Plant Extract on Escherichia coli0157:H7. Pakistan Journal of Nutrition. 5, 325–327.

Aboaba OO, Efuwape BM. 2001. Antibacterial properties of some Nigerian species. Biophysical Research Communications. 13, 183–188.

Adom KK, Sorrells ME, Liu RH. 2005. Phytochemicals and antioxidant activity of milled fractions of different wheat varieties. Journal of Agricultural and Food Chemistry. 53, 2297–2306.

Ammon HP, Muller AB. 1985. Forskolin: from an Ayurvedic remedy to a modern agent. Planta Medica. 6, 473–477.

Awika JM, Rooney LW. 2004. Sorghum phytochemicals and their potential impact on human health. Phytochemistry. 65, 1199–1221.

Aziagba BO, Okeke CU, Ezeabara AC, Ilodibia CV, Ufele AN, Egboka TP. 2017. Determination of the Flavonoid Composition of Seven Varieties of Vigna unguiculata (L.) Walp as Food and Therapeutic Values. Universal Journal of Applied Science. 5(1), 1–4.

Balasundram N, Sundram K, Samman S. 2006. Phenolic compounds in plants and agri industrial by-products: antioxidant activity, occurrence, and potential uses. Food Chemistry. 99, 191–203.

Brunner JH. 1984. Direct spectrophotometric determination of Saponins. Analytical Chemistry. 34, 1314–1326.

Chang C, Yang M, Wen H, Chern J. 2002. Estimation of total flavonoid Content in propolis by two complementary colorimetric methods. Journal of Food and Drug analysis. 10, 178–182.

Chowdhary AR, Sharma ML. 1998. GC-MS investigations on the essential oil from Coleus forskohlii Briq. Indian perfumer. 42, 15–16.

Clarice P, Mudzengi, Amon M, Musa T, Chrispen M, Joan V, Burumu, Tinyiko H. 2017. Antibacterial activity of aqueous and methanol extracts of selected species used in livestock health management. Pharmaceutical Biology. 55(1), 1054–1060.

Cseke LJ, Kirakosyan A, Kaufman PB, Warber S, Duke JA, Brielmann HL. 2016. Natural products from plants. CRC Press.

El-olemy MM, Al-muhtadi FJ, Afifi AFA. 1994. Experimental Phytochemistry. A laboratory manual. 21–27.

Foyer CH, Lelendais M, Kunert KJ. 1994. Photooxidative stress in plants. Physiologia Plantarum. 92, 696–717.

Ghazghazi H, Chedia A, Moufida W, Faten T, Abderrazak M, Brahim H. 2015. Chemical composition of Ruta chalepensis leaves essential oil and variation in biological activities. Journal of Essential Oil Bearing Plants. 18, 3.

Gil A, De La Fuente EB, Lenardis AE, Loopez Pereira M, Suaorez SA, Bandoni A, Van Baren C, Di Leo Lira P, Ghersa CM. 2002. Coriander essential oil composition from two genotypes grown in different environmental conditions. Journal of Agricultural Food Chemistry. 50, 2870–2877.

Hooper DC. 2001. Emerging mechanisms of fluoroquinolone resistance. Emerging Infectious Diseases Journal. 7, 337–341.

Ibrahim SA, Idris AN, Abayomi S, Fatima Y, Auwal AA, Ismail AH. 2015. Phytochemical Screening and Antimicrobial Activities of Ethanolic Extracts of Moringa oleifera Lam on Isolates of Some Pathogens. Journal of Applied Pharmaceutical Science. 7; 4.

Indumathi C, Durgadevi G, Nithyavani S, Gayathri PK. 2014. Estimation of terpenoid content and its antimicrobial property in Enicostemma litorrale. International Journal of Chem Tech Research. 6(9), 4264–4267.

Jacqueline ET, Christian UI. 2017. Evaluation of Anti-fungal Activity of Coleus Species Extracts. International Journal of Current Research in Biosciences and Plant Biology. 4(1), 131–138.

Jagtap NS, Khadabadi SS, Ghorpade DS, Banarase NB, Naphade SS. 2009. Antimicrobial and antifungal activity of Centella asiatica (L.) Urban, Umbeliferae. Research Journal of Pharmacy and Technology. 2(2), 328–330.

Joshi H, Parle M. 2006. Cholinergic basis of memory improving effect of Ocimum tenuiflorum Linn. Indian Journal of Pharmaceutical Science. 68(3), 364–365.

Kate VV. 2008. Physiological and biochemical studies in some medicinal plants: Tribulus terrestris L. and Pedalium murex L. Ph. D. Thesis submitted to Shivaji University, Kolhapur, Maharashtra, India.

Kent T, Kirk, John R, Obst. 1988. Lignin Determination. Methods in Enzymology. 161, 87–110.

Ksouri R, Megdiche V, Debez A, Falleh H, Grignon C, Abdelly C. 2007. Salinity effects on polyphenol content and antioxidant activities in leaves of the halophyte Cakile maritime. Plant Physiology and Biochemistry. 45, 244–249.

Lee JC, Lee KY, Kim J, Na CS, Jung NC, Chung GH, Jang YS. 2004. Extract from Rhus verniciflua stokes is capable of inhibiting the growth of human lymphoma cells. Food and Chemical Toxicology. 42(9), 1383–1388.

Malick CP, Singh MB. 1980. In: Plant Enzymology and Histoenzymology. Kalyani Publishers. 286.

Malleswari D, Mohd KM, Rana K, Bagyanarayana G. 2017. Antibacterial and Antifungal Activity of Leaf, Stem and Root Extracts of Costus Igneus. Research Journal of Pharmaceutical, Biological and Chemical Sciences. 8; 2314.

Maragathavalli S, Brindha S, Kaviyarasi NS, Annadurai BB, Gangwar SK. 2012. Antimicrobial activity in leaf extract of Neem (Azadirachta indica Linn.). International Journal of Science and Nature. 3(1); 110–113.

Marosz A, Nowak J.S. 2008. Effect of salinity stress on growth and macro elements uptake of four tree species. Dendrobiology. 59, 23–29.

Mboso OE, Eyong EU, Odey MO, Osakwe E. 2013. Comparative phytochemical screening of Ereromastax speciosa and Ereromastax polysperma. Journal of Natural Product and Plant Resources. 3, 37–41.

Muthukumarasamy M, Gupta SD, Pannerselvam R. 2000. Enhancement of peroxidase, polyphenol oxidase and superoxide dismutase activities by tridimefon in NaCl stressed Raphanus sativus L. Biology of Plant. 43, 317–320.

Muthumani P, Meera R, Sweetlin, Devi P. 2010. Phyto Chemical Investigation and Determination of Crude Alkaloidal Content (Solasodine) in Solanum Leave Dunal (Dry and Fresh Berries). International Journal of Pharmaceutical & Biological Archives. 1, 350–354.

Nahak G, Sahu RK. 2010. In-vitro antioxidative activity of Azadirachta indica and Melia azedarach leaves by DPPH scavenging assay. Natural Science. 8(4), 22–28.

Naima S, Muhammad RK, Maria S. 2012. Antioxidant activity, total phenolic and total flavonoid contents of whole plant extracts Torilis leptophylla L. BMC Complementary and Alternative Medicine. 12, 221.

Narendra D, Ramalakshmi N, Satyanarayana B, Sudeepthi P, Hemachakradhar K, Pavankumar raju N. 2013. Preliminary Phytochemical Screening, Quantitative estimation and Evaluation of anti-microbial activity of Alstoniamacrophylla Stem bark. International Journal of Science Inventions Today. 2, 31–39.

Navarro JM, Flores P, Garrido C, Martinez V. 2006. Changes in the contents of antioxidant compounds in pepper fruits at ripening stages, as affected by salinity. Food Chemistry. 96, 66–73.

Odeja O, Grace O, Christiana EO, Elias EE, Yemi O. 2015. Phytochemical Screening, Antioxidant and Antimicrobial activities of Senna occidentalis (L.) leaves Extract. Clinical Phytoscience. 1, 6.

Ouda SAE, Mohamed SG, Khalil FA. 2008. Modelling the effect of different stress conditions on maize productivity using yield-stress model. International Journal of Natural and Engineering Sciences. 2, 57–62.

Pandey G, Madhuri S. 2010. Pharmacological activities of Ocimum sanctum (Tulsi): A Review. International Journal of Pharmaceutical Sciences Review and Research. 5(1); 61–66.

Polshettiwar SA, Ganjiwale RO, Wadher SJ, Yeole PG. 2007. Spectrophotometric estimation of total tannins in some Ayurvedic eye drops. Indian Journal of Pharmaceutical Science. 69, 574–576.

Prashant A, Mehta JP. 2017. A review on antimicrobial and Himalayan medicinal plants. Environment Conservation Journal. 18; 49–62.

Ramadoss D, Lakkineni VK, Bose P, Ali S, Annapurna K. 2013. Mitigation of salt stress in wheat seedlings by halo tolerant bacteria isolated from saline habitats. Springer Plus. 2, 1–7.

Sewelam N, Kemel K, Peer MS. 2016. Global plant stress signalling: Reactive oxygen species at the cross-road. Frontiers of Plant Science. 7, 187.

Seyyednejad SM, Motamedi H. 2010. A review on native medicinal plants in Khuzestan, Iran with antibacterial properties. International Journal of Pharmaceutics. 6, 551–60.

Sharma A, Meena A, Meena R. 2012. Antimicrobial activity of plant extract of Ocimum tenuiflorum. International Journal of Pharma Tech Research. 4(1); 176–180.

Shuji Y, Ray AB, Paul MH. 2002. Salt stress tolerance of plants. JIRCAS Working Report. 25–33.

Soladoye MO, Chukwuma EC. 2012. Quantitative phytochemical profile of the leaves of Cissus populnea Guill. & Perr. (Vitaceae) – An important medicinal plant in central Nigeria. Scholars Research Library. Archives of Applied Science Research. 4, 200–206.

Surh YZ, Ferguson LR. 2003. Dietary and medicinal antimutagens and anticarcinogens. Molecular mechanisms and chemopreventive potential-highlight of a symposium.

Taylor JR. 1982. An Introduction to Error Analysis. The Study of Uncertainties in Physical Measurements. University Science Books, Sausalito, CA, USA.

Tester M, Davenport R. 2003. Na+ tolerant and Na+ transport in higher plants. Annals of Botany. 91, 503–527.

Thilagavathi S, Hariram N. 2016. Coleus aromaticus Benth Synthesis of Potentially Nanomedicine as High Nutritive Value of Human Health and Immunomodulator. International Journal of Science and Research Methodology. 4(4); 18–38.

Thuy TV, Hyungrok K, Vu KT, Quang LD, Hoa TN, Hun K, In Seon K, Gyung JC, Jin-Cheol K. 2016. In vitro antibacterial activity of selected medicinal plants traditionally used in Vietnam against human pathogenic bacteria. BMC Complementary and Alternative Medicine. 16; 32.

Valifard M, Mohsenzadeh S, Kholdebarina B, Rowshanb V. 2014. Effects of salt stress on volatile compounds, total phenolic content and antioxidant activities of Salvia mirzayanii. South African Journal of Botany. 93, 92–97.

Vimala G, Francis GS. 2015. Qualitative and quantitative determination of secondary metabolites and antioxidant potential of ficus benghalensis linn seed. International Journal of Pharmacy and Pharmaceutical Sciences. 7(7).

Vimala V, Rebecca Mathew, P, Deepa S, Kalaivani T. 2014. Phytochemical analysis in ocimum accessions. International Journal of Pharmacy and Pharmaceutical Sciences. 6(1).

Warrier PK, Nambiar VP, Ramankutty C. 1995. Indian Medicinal Plants. Orient Longman Ltd., Madras.

William PCB, Raquel OR, Demetrio LV, Juliana JMP. 2016. Bioactive metabolite profiles and antimicrobial activity of ethanolic extracts from Muntingia calabura L. leaves and stems. Asian Pacific Journal of Tropical Biomedicine. 6(8), 682–685.

Wong CC, Li HB, Cheng KW, Chen F. 2006. A systematic survey of antioxidant activity of 30 Chinese medicinal plants using the ferric reducing antioxidant power assay. Food Chemistry. 97, 705–711.

Xin ZL, Mei G, Shiqing L, Shengxiu L, Zongsuo L. 2011. Modulation of plant growth, water status and antioxidative system of two maize (Zea mays L.) cultivars induced by exogenous glycinebetaine under long term mild drought stress. Pakistan Journal of Botany. 43, 1587–1594.

Yala D, Merad AS, Mohamedi D, Ouar Korich MN. 2001. Classification et mode d’action des antibiotiques. Médecine du Maghreb. 91, 5–12.

